# Unraveling phenotypic variation in multiscale behavioral dynamics

**DOI:** 10.64898/2026.10.05.756657

**Authors:** Gautam Sridhar, Claire Wyart, Antonio Carlos Costa

## Abstract

Behavioral variability reflects differences in internal and external parameters and provides a substrate for natural selection. However, uncovering its structure is challenging because behavior evolves nonlinearly across multiple timescales. Our goal is to reconstruct phenotypic variation from finite behavioral observations without prescribing the scales at which individuals should be compared. To achieve this, we introduce a multiscale dissimilarity that compares dynamics across all statistically resolvable resolutions by constructing reduced transfer operators at progressively finer scales and weighting their differences by finite-sampling uncertainty. This dissimilarity defines a geometry of dynamical phenotypes, which we reconstruct using diffusion maps. We show that the resulting phenotypic coordinates locally follow the parameter directions to which dynamics are most sensitive. We validate our approach in a stochastic model and apply it to bacterial and larval zebrafish behavior. In bacteria, we identify run speed as a major component of inter-individual variability, associated with the chemotaxis protein CheB. In zebrafish, we find that prior prey experience reshapes behavioral dynamics and phenotypic variability, with paramecia experience concentrating fish in a prey-capture-associated region of phenotypic space. Together, these results provide a framework for identifying internal and environmental variables that structure dynamical phenotypes across scales.

## INTRODUCTION

Behavior at the scale of an entire organism is uniquely challenging to study as it operates across multiple spatial and temporal scales with substantial variability, complicating efforts to extract general principles. A common approach to cope with this variability is to average it away - in neuroscience, for example, averaging measurements to discover significant trends or effects of manipulations is widespread [1]. Yet, the variation across realizations may itself carry biological signal, reflecting differences in internal state [2] that shape how individuals respond variably to the same input [3]. Phenotypic variation also constitutes the substrate for natural selection: because different behavioral strategies are optimal in different ecological contexts, the spread of phenotypes in a population can shape a population’s capacity to adapt to changing environments [4, 5]. Variability must therefore be treated not as noise to be averaged away, but as a signal reflecting the constraints that shape the parameters underlying the emergent dynamics, Fig. 1(a1). Our goal is to reconstruct the structure of behavioral variation from finite observations, identify the dynamical scales at which individuals differ, and relate this variation to the internal and environmental variables that shape it.

**FIG 1.**
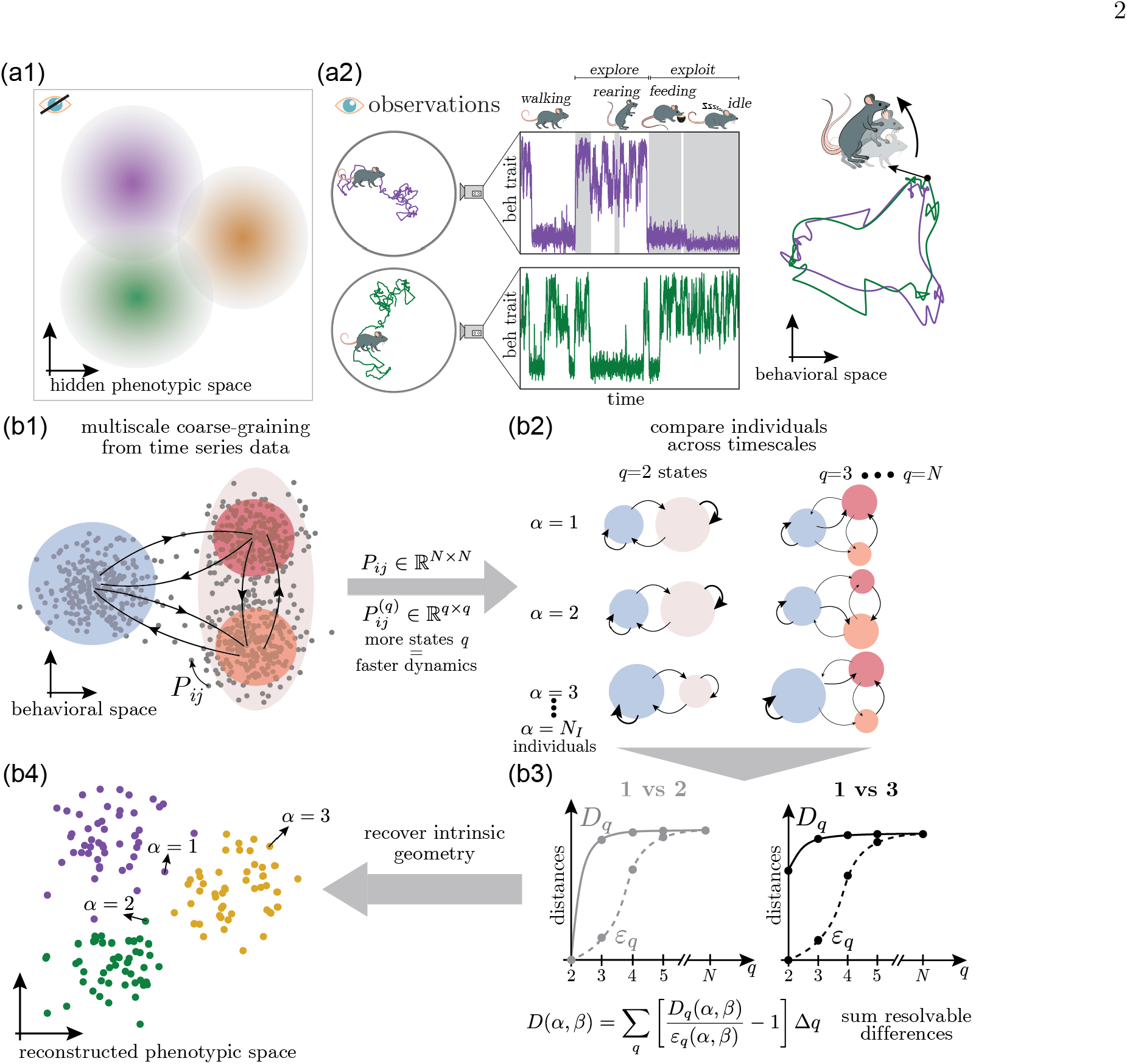
Unraveling behavioral variability across timescales. (a) We aim to reconstruct hidden behavioral phenotypes (a1) from behavioral observations (a2). Quantitative approaches enable the construction of rich behavioral representations [10–12]. However, quantifying inter-individual differences remains a challenge as differences can manifest across timescales. For example, two individuals may use the same short time motifs, but sequence them in different ways over time. (b) Our approach compares these dynamics without selecting a coarse-graining scale *a priori*. (b1) Starting from a fine scale Markov model *P*_*ij*_ ∈ R^*N*×*N*^ among *N* short-time movements, we identify *q* metastable states and construct reduced-order *q*-state models, 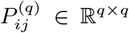, with an increasing number of states *q* capturing faster dynamics. (b2) Each individual trajectory is projected onto the resulting family of Markov models. (b3) At each resolution *q*, we measure the dynamical difference between individuals *D*_*q*_ (*α, β*), Eq. 1, and compare it with a finite-sampling uncertainty scale *ε*_*q*_ (*α, β*), Eq. 2. This enables the quantification of dynamical differences across time scales. For example, individuals 1 and 2 share the same dynamics at the level of *q* = 2 states, but differ in the finer scale transition at *q* = 3. In contrast, individual 3 already differs from individual 1 at the level of *q* = 2 states. As *q* increases, sampling uncertainty eventually makes finer differences unresolvable. We sum the excess of *D*_*q*_ (*α, β*) over this uncertainty *ε*_*q*_ (*α, β*) across all resolvable scales to obtain a single multiscale dissimilarity, *D*(*α, β*), Eq. 3. (b4) Diffusion maps reconstruct the intrinsic geometry induced by these pairwise dissimilarities, yielding a phenotypic space that reflects significant dynamical differences across scales.

While relatively static phenotypic differences can be quantified through precise measurements [6], phenotypes that are dynamic in nature, such as behavior or neural dynamics, are harder to compare. Common approaches rely on time-averaged quantities, such as mean speed or the frequency of discrete behaviors [7–9]. While insightful, these approaches miss the dynamic and multiscale nature of behavior [10–12], Fig. 1(a2). Two individuals may, for example, have similar exploration vs. exploitation propensities, but vary in the finer scale movements used throughout. Comparing dynamical phenotypes therefore requires a representation that can resolve differences across scales without prescribing in advance which scales matter.

To achieve this, we construct a dissimilarity between finite dynamical observations that compares coarse-grained dynamics across a range of resolutions, Fig. 1(b). At each resolution, observed differences are calibrated against the uncertainty expected from finite sampling, so that only empirically resolvable dynamical structure contributes to the comparison. The resulting dissimilarity defines a nonlinear geometry of dynamical phenotypes, which we reconstruct using diffusion maps. We further show that the resulting coordinates locally follow parameter directions to which the dynamics are most sensitive (see Supplementary Information).

We illustrate the method in a stochastic model and apply it to bacterial and larval zebrafish behavior. In bacterial motility, we find that differences in run speed are a major component of inter-individual variation and are strongly associated with variation in CheB expression. In zebrafish, we find that prior prey experience reshapes both behavioral strategies and the distribution of phenotypes in the population, with distinct effects for different prey types.

## RESULTS

### Reconstructing phenotypic geometry from multiscale dynamics

To compare dynamics across timescales, we build upon recent work [13–16] and approximate the dynamics through discrete versions of linear transfer operators *P*_*ij*_, which can successfully encode nonlinear dynamics [17]. Given a set behavioral observations, we first construct an ensemble transition matrix *P*_*ij*_ across all the realizations to obtain a common behavioral space, Fig. 1(b1) (see Methods). Its non-trivial eigenvectors identify long-lived dynamical structure and provide a basis for coarse-graining into metastable states [18–20] (see Methods). Using increasingly many eigenvectors, we obtain a family of *q*-state Markov models 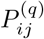, with larger *q* resolving progressively faster dynamics, Fig. 1(b1,b2).

For each coarse-graining scale *q*, we measure the difference between realization *α* and *β* using the average total-variation distance among the rows of their coarse-grained transition matrices,

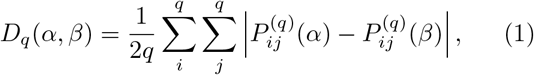

obtaining a measure of the difference between the dynamics at a specific timescale set by *q* (the factor of 2 bounds the distance 0 ≤ *D*_*q*_(*α, β*) ≤ 1), Fig. 1(b3).

Crucially, we then determine which differences across multiple scales *q* are actually resolvable from finite data. As *q* increases, trajectories are divided among more states and fewer transitions are available to estimate each element of *P* ^(*q*)^. Consequently, two symbolic sequences generated by the same underlying Markov model can exhibit an apparent distance. We quantify this finite-sampling contribution by resampling under the null hypothesis that both observations were generated by the same *q*-state Markov model and defining the uncertainty scale,

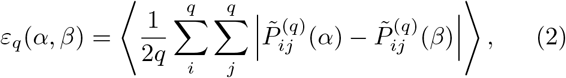

where 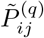 represents the resampled transition matrices and the expectation value is taken across bootstrapped resamples (see Methods).

Beyond some resolution *q*, the measured distance between two individuals is no longer distinguishable from what sampling noise alone would produce, and *D*_*q*_(*α, β*) *→ ε* _*q*_(*α, β*). We therefore combine scales by accumulating, across *q*, the amount by which the measured distance exceeds this uncertainty scale,

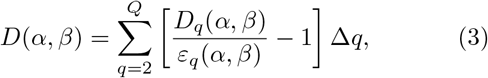

where *Q* is taken large enough so that *D*_*q*_*/ε*_*q*_ ≈1. The resulting dissimilarity thus combines all empirically resolvable dynamical differences without selecting a preferred coarse-graining in advance.

Given these dissimilarities, we next ask how phenotypic variation is structured. Individuals differ in many underlying interacting parameters (e.g. protein expression levels, neuromodulatory states) but these need not map onto dynamical differences in a simple way [21]: the same change in a parameter can have a large effect in one region of parameter space and none in another. In addition, evolutionary trade-offs may confine phenotypes to a low-dimensional, nonlinear manifold [22, 23], frequently with a simplex structure [6, 24, 25]. We therefore expect phenotypic variation to occupy a potentially curved, low-dimensional space whose geometry reflects how parameter variation is expressed in dynamics. To reconstruct this geometry from the pairwise dissimilarities *D*(*α, β*), we use diffusion maps [26], which recover the intrinsic geometry of a set of points by constructing a random walk that hops between nearby points, Fig. 1(b4). We use a self-tuning kernel bandwidth, so that the walk adapts to the local density of phenotypes, and a density normalization ensuring that the recovered geometry reflects the structure of the manifold rather than how densely individuals happen to sample it (see Methods).

This choice is not merely practical. We show that the diffusion coordinates recovered in this way locally align with the dominant modes of the metric tensor mapping parameter variation onto emergent differences in dynamics (see Supplementary Information): the directions along which the embedding separates individuals most strongly are the directions in parameter space to which the dynamics are most sensitive. The geometry we reconstruct is therefore not an arbitrary visualization of the distances, but a representation of how parameter variation is expressed in behavior. Our approach thus constructs phenotypic spaces that respect the multiscale nature of behavior, weight differences by their significance, and reflect the nontrivial relationship between underlying parameters and observed dynamics. Below we illustrate each step of our approach in a toy model.

#### Multiscale total variation dissimilarity: a working example

We consider overdamped dynamics in a triple-well potential, varying the depths of the three minima, *µ*_1_, *µ*_2_, *µ*_3_, to generate four dynamical “phenotypes”. Parameter values vary weakly within each group and more strongly across groups, producing correspondingly larger differences in the emergent dynamics. We further constrain the underlying parameter variation to a two-dimensional plane, Fig. 2(a), so that the geometry to be reconstructed is known. This model allows us to test whether the multiscale dissimilarity identifies the resolution at which dynamical differences emerge, suppresses structure that cannot be resolved from finite data, and recovers the known low-dimensional organization of the underlying parameters.

**FIG 2.**
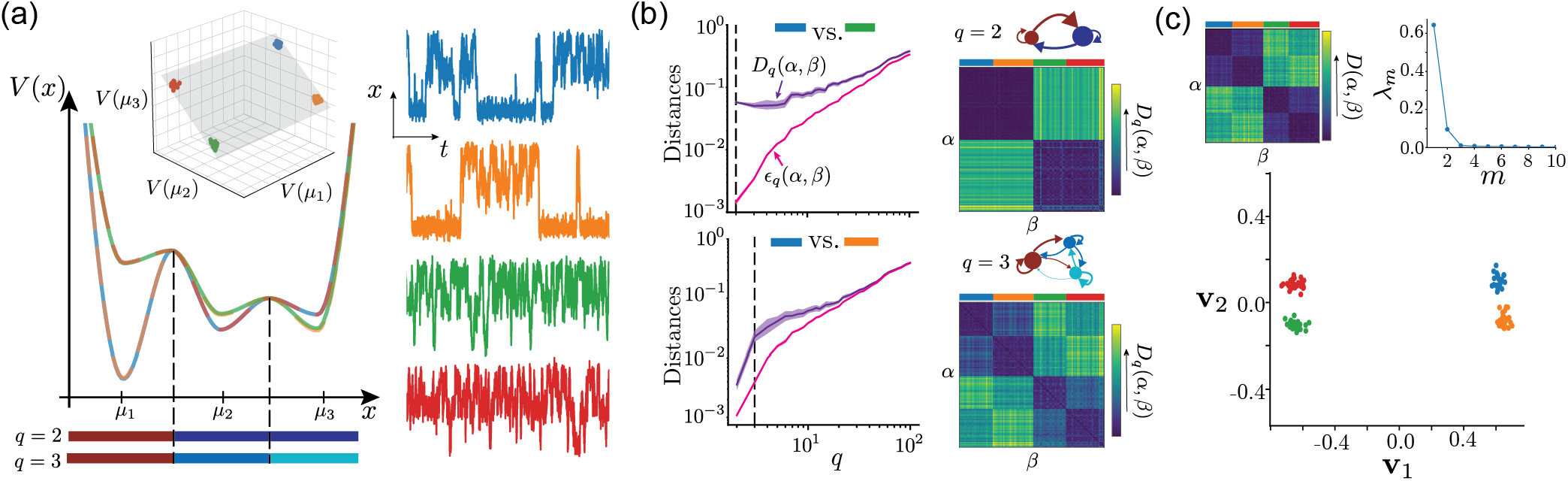
Reconstructing the phenotypic space of variation among overdamped stochastic dynamics in a 3-well potential. (a) We simulate overdamped Langevin dynamics in a potential with minima at *µ*_1_, *µ*_2_, *µ*_3_, and generate four dynamical “phenotypes” by varying the depths of the three wells. We couple *V* (*µ*_2_) and *V* (*µ*_3_) so that the underlying parameter variation is confined to a two-dimensional plane (inset), and show representative 200 s trajectories from each phenotype (right). The colored bars below the potential indicate the coarse-grained states inferred from the dynamics: at *q* = 2, the highest energy barrier separates the system into two metastable states, whereas at *q* = 3 the shallower barrier is additionally resolved. (b) Scale-dependent dissimilarities *D*_*q*_ (*α, β*) reveal the resolution at which phenotypic differences emerge. We compare the blue phenotype with the green (top) and orange (bottom) phenotypes. Blue and green are already well separated at *q* = 2, whereas distinguishing blue from orange requires the finer *q* = 3 description, which resolves transitions between the two shallower wells. The corresponding coarse-grained Markov models and pairwise distance matrices for *q* = 2 and *q* = 3 are shown on the right. The finite-sampling uncertainty *ε*_*q*_ (*α, β*), Eq. 2, increases with *q* and approaches *D*_*q*_ for large *q*, beyond which additional differences are no longer statistically resolvable. These scale-dependent quantities are combined into the multiscale dissimilarity *D*(*α, β*), Eq. 3. (c) The resulting multiscale dissimilarity matrix (top left) retains the separation among the four phenotypes. Its diffusion-map spectrum (top right) exhibits two dominant non-trivial modes, consistent with the two-dimensional parameter variation imposed in (a). Projection onto these two diffusion coordinates (bottom) recovers the known low-dimensional organization and separates all four dynamical phenotypes.

The coarse-grained models resolve the metastable structure progressively as *q* increases, Fig. S1 (see Methods). The leading non-trivial eigenvector of the reversibilized transfer operator, *ϕ*_1_, partitions the dynamics into *q* = 2 metastable states separated by the highest energy barrier, Fig. S1(b). Including the second eigenvector, *ϕ*_2_, yields a *q* = 3 description that additionally resolves transitions across the lower barrier, Fig. S1(c). For *q>* 3, the coarse-graining further subdivides transition regions while preserving the three metastable basins, Fig. S1(d).

This scale dependence is reflected directly in the pairwise distances across *q*, Fig. 2(b). Blue and green phenotypes are already well separated at *q* = 2, whereas properly distinguishing blue from orange requires the finer *q* = 3 description. Beyond *q* = 3, no additional metastable structure separates the phenotypes, and finer subdivisions capturing differences in the local curvature of the potential landscape contribute little to *D*_*q>*3_. At the same time, the finite-sampling uncertainty *ε*_*q*_ increases with *q* and approaches *D*_*q*_ at sufficiently fine resolution. The multiscale dissimilarity, Eq. 3, therefore retains the resolutions at which phenotypes differ while progressively suppressing differences that become indistinguishable from sampling fluctuations. Consistently, shortening the observation time causes fine-scale differences to lose significance earlier, Fig. S2.

Finally, diffusion maps recover two dominant non-trivial coordinates, matching the two-dimensional parameter variation imposed in Fig. 2(a). The leading coordinate **v**_1_ captures the larger differences associated with the left well, while **v**_2_ resolves the smaller differences within the right basin, separating all four phenotypic groups, Fig. 2(c). Thus, the toy model shows that the proposed dissimilarity retains dynamical differences at the resolutions supported by the data and recovers their underlying low-dimensional organization.

### Applications to data

We now consider real-world datasets for which the structure of phenotypic variation may reflect variation in measurable internal mechanisms or external conditions.

#### From gene expression to behavioral variation in E. Coli motility

The motility of *E. coli* provides a useful test case because its chemotaxis network is well characterized, while substantial phenotypic variation persists even within isogenic populations [29, 30] and has been linked to adaptability [5, 27]. We analyze the dataset of Dufour et al. [27], which combines motility trajectories from wild-type cells with measurements from strains in which the expression of the adaptation proteins CheR and CheB was controlled and quantified, Fig. 3(a). CheR and CheB regulate receptor methylation and thereby influence CheY-P activity and tumble bias. We ask whether phenotypic variation extends beyond the canonical descriptors of mean run speed and tumble bias, and how the resulting dynamical variation relates to CheR and CheB abundance.

**FIG 3.**
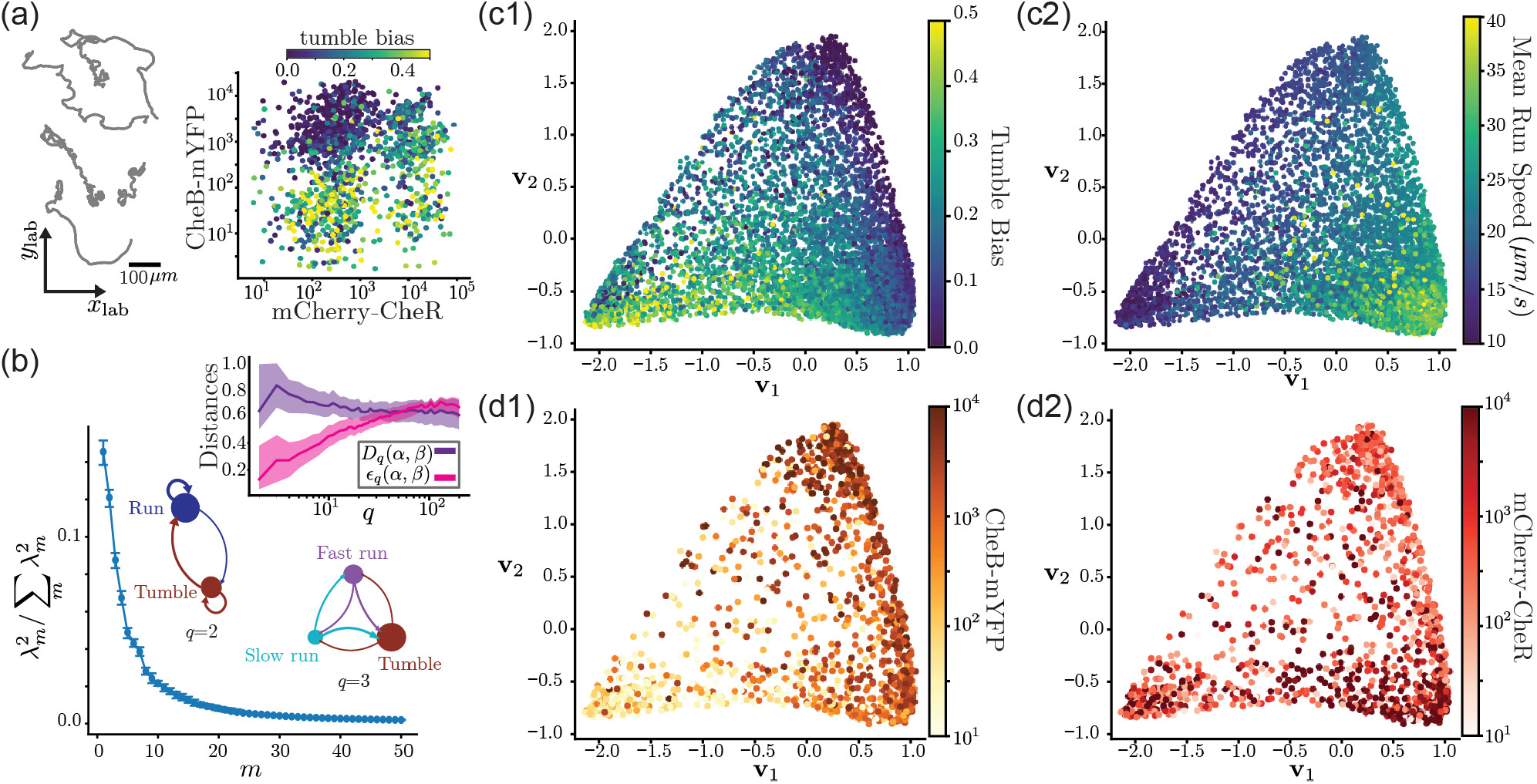
Phenotypic space of *E. coli* motility highlights the importance of speed modulation, and the primary impact of CheB on inter-individual variability. (a-left) Example trajectories from the dataset [27], which consists of RP437 isogenic wild-type bacteria (5399 tracks) and two mutant strains (1374 tracks) modified to control the expression of CheR and CheB. Trajectories last an average of 48.4 (10.8, 191.6) s. (a-right) CheR and CheB numbers for each mutant cells, color-coded by the tumble bias (as in Fig.3D of [27]). High values of CheB and low values of CheR correspond to low tumble bias (indicative of straight runs). High values of CheR when CheB numbers are high increases the tumble bias. Low CheB values display overall higher tumble bias irrespective of variation in CheR. (b) The kinetic variance [28] captured by the eigenspectrum of the diffusion map reveals a structured phenotypic space where a few coordinates capture most of the kinetic variance. (b-inset) Distances *D*_*q*_ and uncertainties *ε*_*q*_ between coarse-grained operators *P* ^(*q*)^. The differences peak at *q* = 3 and are resolved by *q ≈* 20, after which the difference in phenotypes become statistically insignificant. We also show the *q* = 2 and *q* = 3 transition matrices for an example cell, showcasing how at *q* = 3 the dynamics incorporate slower and faster runs, which are important for capturing inter-individual differences. (c) We plot the position of each cell along the two major axis of phenotypic variation, and color code them by tumble bias (c1) and mean run speed (c2). We find important difference in tumble bias along both **v**_1_ and **v**_2_, whereas variation is run speed occurs mostly along **v**_1_ only, see Fig. S4 for details. (d) Focusing on mutant strains, we find that most of the phenoypic variation aligns with differences in CheB, see also Fig. S4(a). Both **v**_1_ and **v**_2_ are correlated with differences in CheB (d1), with **v**_2_ also showing some negative correlations with CheR when **v**_1_ is large (d2). This indicates that variation in CheB numbers has a primary impact on the phenotypic variability, whereas differences in CheR only becomes relevant when CheB numbers are su”ciently large.

To reconstruct the phenotypic space, we start by inferring the long-lived dynamics from the observed center-of-mass trajectories, see Fig. S3 and Methods for details. We directly uncover two timescale separated behavioral modes. The slowest mode is the canonical run-and-tumble [31, 32], with ≈ 2*s* mixing time. The average duration of tracks is 48.4 (10.8, 191.6) s, and so the tumble bias is approximately constant [33] and constitutes the slowest timescale in the observed dynamics. The second mode differentiates fast and slow runs (≈1*s* average mixing time). Beyond this, other modes have negligible timescale separation but are still interpretable: for example, the third and fourth modes capture differences in the straightness and left-right asymmetry of trajectories, Fig. S3, which may however arise due to boundary effects. Overall, the *q* = 2 states correspond to runs and tumble, at *q* = 3 we find fast and slow runs vs tumbles, and beyond that we get finer subdivisions of these states that captures curvature and handedness differences.

Next, we proceed to construct our multiscale dissimilarity, as described above. We find that most of the kinetic variance is captured in the first few modes of the diffusion map, Fig. 3(b), pointing to structured variation. The multiscale dissimilarity is concentrated at low *q*, with its strongest contribution around *q* = 3, and becomes indistinguishable from finite-sampling uncertainty by *q ≈*20, Fig. 3(b). Because the *q* = 3 representation specifically resolves fast and slow runs, this peak indicates that variation in run-speed dynamics contributes substantially to differences among cells, beyond tumble bias alone, mirroring what has been found in [33].

Projecting cells onto the two dominant diffusion coordinates reveals broad correspondence with canonical behavioral statistics, but not a complete reduction to them, Fig. 3(c). A linear regression of the position along **v**_1_ and **v**_2_ from speed and tumble bias yielding an *R*^2^ = 0.555 (0.546, 0.567) (where the parenthesis represent bootstrapped 95% confidence intervals), Fig. S4(a). Nonetheless, **v**_1_ is strongly correlated with mean run speed, while variation along **v**_2_ is associated with both run speed and tumble bias. Run speeds are also more important than tumble bias in predicting the position of each individual cell in the phenotypic space, Fig. S4(a1-right), indicating that run-speed modulation is a major component of inter-individual variation that is not captured by a description based on tumble bias alone.

We next asked how this phenotypic geometry relates to variation in the chemotaxis network. Among mutant cells, position in the recovered space is more strongly associated with CheB than with CheR abundance, Fig. 3(d),Fig;S4(a2). Both dominant diffusion coordinates vary systematically with CheB, whereas associations with CheR are weaker and become most apparent at high CheB abundance. This asymmetry is consistent with earlier observations in the same dataset [27], but the dynamical decomposition further shows that the dominant CheB-associated axis is also strongly related to run speed. Direct comparison of protein abundance with conventional behavioral statistics confirms that run speed varies systematically with CheB, in addition to the expected associations of CheR and CheB with tumble bias, Fig. S4(a3).

Wild-type and mutant cells also occupy the phenotypic space differently, Fig. S4(b1). The mutant population is more variable and anisotropic, with its dominant variation extending along directions associated with both tumble bias and run speed, Fig. S4(b2). Wildtype variation is more constrained and isotropic, with its major axis being strongly associated with run speed, Fig. S4(b3). Thus, the directions explored by perturbing CheR and CheB do not simply reproduce the dominant variation present in the wild-type population, suggesting that additional sources of variation contribute to wildtype motility phenotypes.

We also noted that the projections along (**v**_1_, **v**_2_) are mostly distributed inside a simplex, prompting an analysis of the differences in behavior in the extremes of the distribution, Fig. 4(a1). We clustered the space to identify three groups (see Methods), and found that these groups differ both in protein abundance and primarily in their *q* = 3 dynamics: *G*_3_ has low CheB and a high tumble propensity, whereas *G*_1_ (high CheB, mostly high CheR) and *G*_2_ (high CheB, low CheR) are distinguished primarily by their use of fast versus slow runs, respectively, Fig. 4(a2,b). We therefore asked whether these dynamical differences have consequences for search performance, Fig. 4(c). Simulations of resource encounter across spatial scales show that *G*_3_ performs best over short distances, whereas *G*_2_ performs best only at much larger scales. *G*_1_ maintains comparatively high performance across a broader range of scales and contains the largest fraction of wild-type cells, Fig. 4(c). This suggests that the average wild-type phenotype is tuned to maintain efficient exploration at multiple length scales, in contrast with the mutant strains which become specialized to either local or global search.

**FIG 4.**
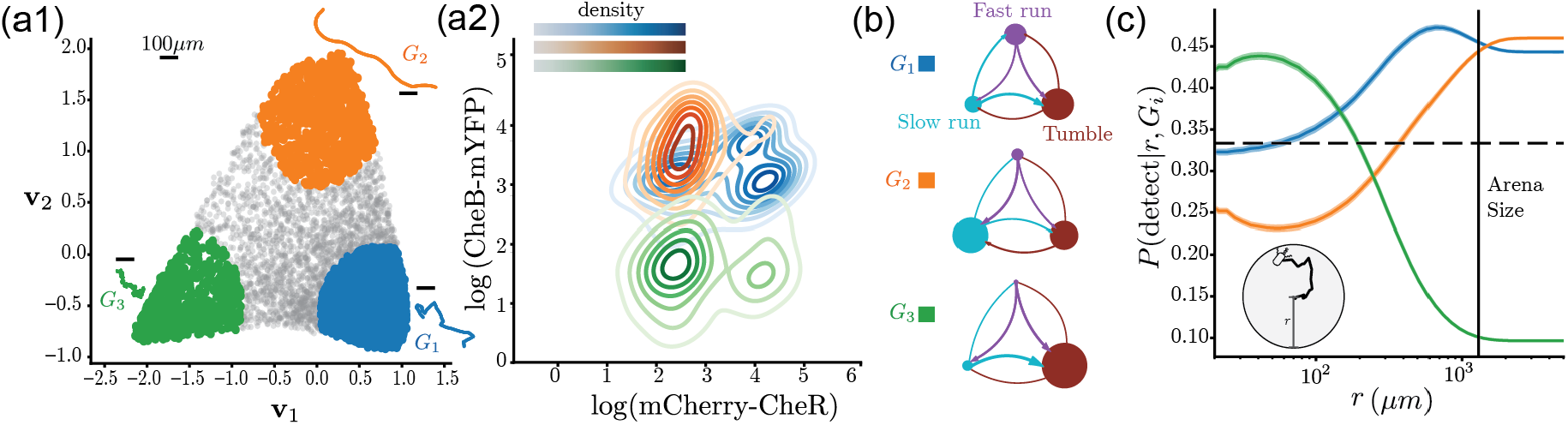
Speed modulation is important for the effectiveness of search strategies. (a1) We analyze the 3 extremes of the phenotypic space in (**v**_1_, **v**_2_), defined using fuzzy c-means (see Methods). We plot example trajectories from each phenotypic group. (a2) Distributions of CheR and CheB numbers for the mutant cells in each of the phenotypic groups. *G*_1_ (blue) exhibits high CheB and variable CheR numbers, *G*_2_ (orange) has high CheB and low CheR numbers and *G*_3_ (green) has low CheB numbers and mostly low CheR numbers. (b) Average *q* = 3 transition matrices in each group. Group *G*_1_ (75.9% wild-type, 24.1% fluorescent) showcases a bias for fast runs and tumbling, group *G*_2_ (21.4% wild-type, 78.6% fluorescent) exhibits more slow runs, and *G*_3_ (19.3% wild-type, 80.7% fluorescent) showcases high bias for tumbling. (c) The probability to detect uniformly distributed resources in a circle of radius *r*. We find that *G*_3_ is most efficient at short length scales, while *G*_2_ only at very large length scales (beyond the arena size of ≈ 10^3^*µ*m). These two groups also have the majority of the mutant bacteria. In contrast, *G*_1_ displays close to average search efficiency at short length scales, is most efficient at length scales comparable to the arena size, and still near optimal at large length scales. This is also the group that includes the most wild-type bacteria. The 1*/*3 horizontal line is the average probability over the 3 clusters.

Altogether, our approach confirmed previous findings on the importance of tumble bias and how it is controlled by CheB. In addition, we also highlighted the importance of run speeds in phenotypic variation and how it relates to CheB expression. Finally, we also found that variability in wild-type cells is not directly aligned the phenotypic axis along which CheR and CheB vary, likely reflecting the tight control of CheR and CheB numbers in wild-type cells resulting in efficient search across length scales.

### Larval zebrafish navigation: multiscale variation highlights impact of prey exposure and developmental history

To further illustrate the potential of our approach at resolving inter-individual variability, we move from bacteria to vertebrate behavior, and leverage a published dataset of larval zebrafish exposed to a variety of sensory contexts, such as aversive visual stimuli or prey of different types [16, 34]. In earlier work [16], we used an approach similar to the one used here for uncovering a hierarchy of predictive Markov models for zebrafish behavior. However, for studying inter-individual variability we resorted to a fixed behavioral representation, which was defined by identifying a set number of coarse-grained states that could best classify all sensory contexts, which turned out to be *q* = 7 states. Here, the comparison is instead performed across all resolvable coarse-graining scales, with each resolution weighted by whether its differences exceed finite-sampling uncertainty.

In this dataset, each recording consists of a sequence of bouts, which results in distinct macroscopic movement patterns. Examining the two example behavioral sequences in Fig. 5(a) highlights the multiscale structure of the behavior that challenges the study of inter-individual variation. Working from bout sequences, we proceed to reconstruct the phenotypic space using the multiscale dissimilarity measure introduced above. Our coarse-graining approach recovers the behavioral structure found in [16]: from cruising and wandering, to fast and slow variations, to left-right asymmetries, and so on.

**FIG 5.**
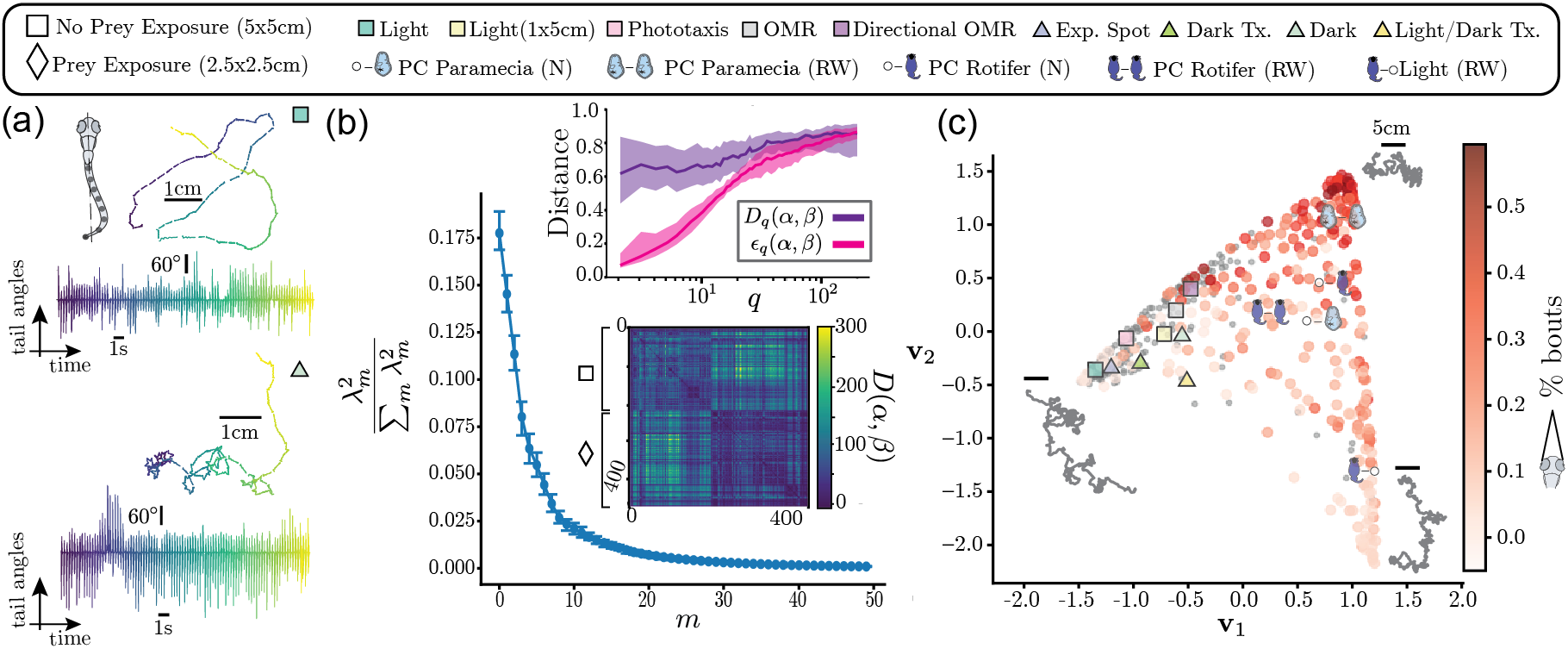
Phenotypic space for larval zebrafish navigation recovers prey exposure as a major determinant of variability. We use the dataset from [16, 34], where 463 larval zebrafish are exposed to different sensory environments (with different tick marks and colors labelled at the top). (a) We showcase two examples of 100 bout sequences and the corresponding fish trajectories, one from fish in the light (top) and one from fish in the dark (bottom). (b) The kinetic variance captured by the eigenvalue spectrum of the diffusion map reveals a structured phenotypic space. (inset, top) Distances *D*_*q*_ (purple) and uncertainties ε_*q*_ (magenta) as a function of coarse-graining scale *q*. On average, across all sensory conditions, there is significant behavioral information up till *q* 20. (inset, bottom) Multiscale dissimilarity, *D*(*α, β*), organized by fish not exposed to prey (square) and fish exposed to prey (diamond). (c) Projection along the first two modes of the diffusion map. Fish in hunting assays are color coded by eye convergence, a proxy for prey capture [35]. We mark the median position in each condition with a special tick mark. The major axis of variation align with the difference between fish that were exposed to prey vs. not, as in [16]. Eye convergence is the highest in the top corner of the phenotypic space.

We find important differences among fish up to *q ≈*20 states, which is well beyond the *q* = 7 state characterization used previously [16]. We sum all significant differences across scales, and obtain a structured phenotypic space with a few diffusion map modes capturing most of the variation, Fig. 5(b). We project all fish along the first two dominant diffusion coordinates, Fig. 5(c). As expected from our previous finding [16], but this time without pre-selecting a specific coarse-graining scale, the first axis neatly separates fish never exposed to prey (negative values) and fish exposed to prey, both prior to or during the experiment (positive values). Low values of **v**_1_ correspond to increased cruising behavior with high speeds [16]. At high **v**_1_, fish exhibit two distinct behavioral patterns depending on **v**_2_: predominantly slow cruising and wandering with high eye convergence (a proxy for prey capture attempts) at high **v**_2_, or predominantly fast wandering with lower eye convergence at low **v**_2_.

As we found in bacteria, the first two diffusion map coordinates **v**_1_ and **v**_2_ form a simplex-like phenotypic space. We therefore examined how fish in the three extremes vary in their search and prey capture abilities (see Methods), Fig. S5. Fish in the left corner are efficient at long distance exploration. Fish in the top corner efficiently search locally and capture prey directly in front of them, whereas fish in the bottom corner are effective at local search but lack frontal-capture ability. Thus, the extremes of the phenotypic space serves either the purpose of local vs global search, or efficiency at capturing prey.

To illustrate what can be learned from the full multiscale representation of zebrafish behavior, we focus on the two conditions with the largest sample sizes in the dataset: in both conditions fish are exposed the paramecia, but in theffnaive” condition fish were not previously exposed to prey (*n* = 65), whereas in the “raised with” condition fish had previously been exposed to prey (*n* = 83). Previous work has shown that prior prey experience strengthens prey-associated visual drive to the forebrain, increasing prey capture attempts and success by tuning the first turn to prey and the speed of the capture maneuver [37, 38]. However, these studies did not resolve how prey exposure can alter behavior on the experimental timescale, neither how it shapes the degree of inter-individual variation among fish.

Our approach uncovers that prior exposure to paramecia reduces inter-individual differences, with a decrease in phenotypic space entropy of *H*_paramecia rw_ −*H*_paramecia naive_ = −1 (−1.5, −0.5) bits, p-value *<* 0.0005 (see Methods), Fig. 6(a). Fish raised with paramecia concentrate at the region of phenotypic space where fish deploy strategies that lead to efficient frontal prey capture, Fig. S5.

**FIG 6.**
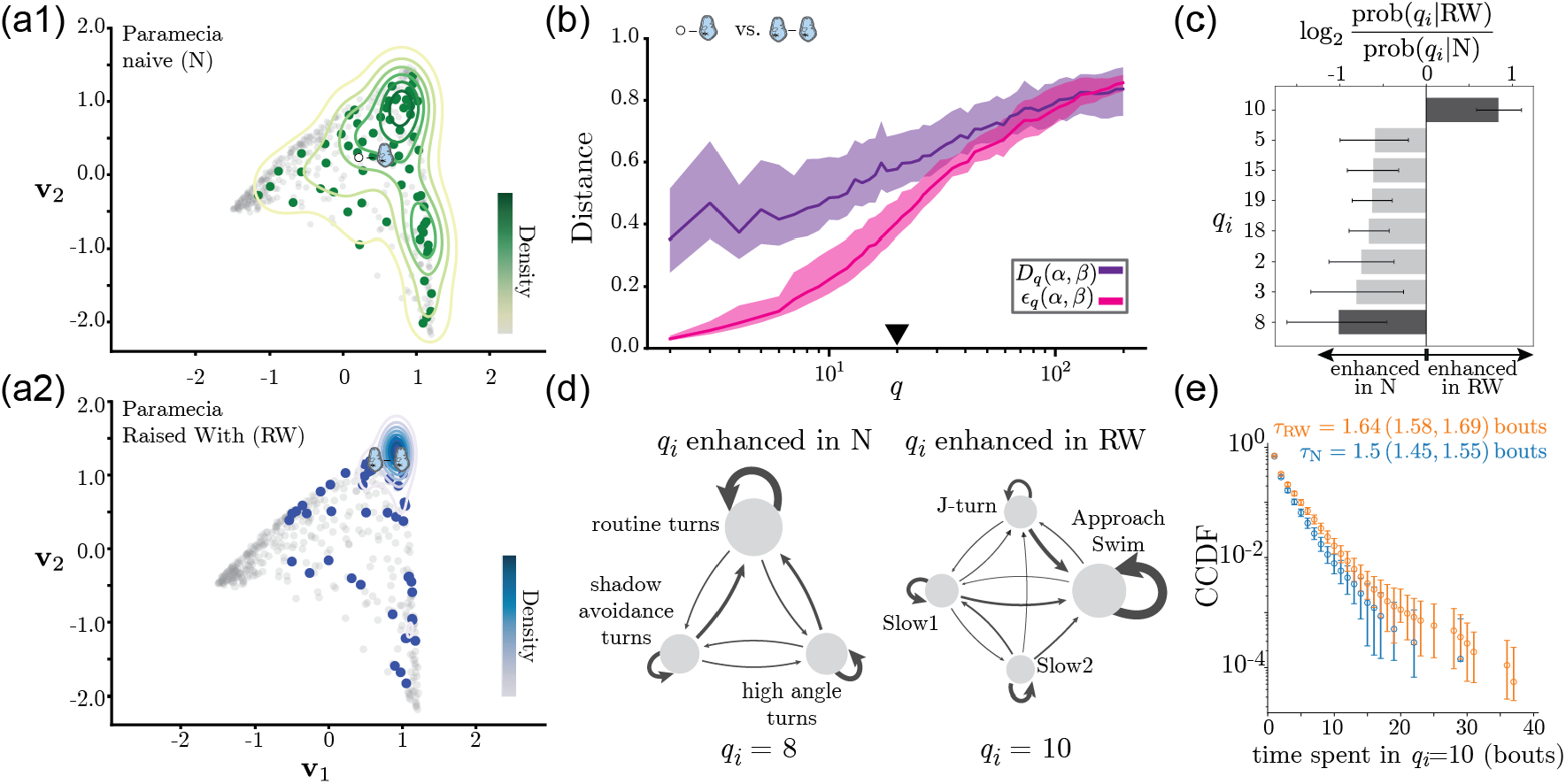
Prior exposure to paramecia reduces inter-individual differences and enhances prey capture sequences. (a1) Density plot of the fish hunting paramecia for the first time. (a2) Density plot of the fish hunting paramecia that were also previously raised with paramecia. Raising the fish with paramecia results in a reduction of inter-individual variability, with the distribution becoming constrained to the upper part of the phenotypic space where eye convergence is high and prey capture is most efficient, Fig. S5. (b) Distances *D*_*q*_ and uncertainty scales *ε*_*q*_ across *q* for comparisons between naive and experience fish hunting paramecia. We find behavioral differences up till *q ≈*20. (c) Log-fold change in the usage of each of the *q*_*i*_ states for *q* = 20. We report only states for which the distribution of bout usage was significantly differ among the two conditions (with p-value *<* 0.01 in a two-sided permutation test and 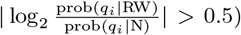, and highlight the two states plotted in panel (d). (d) We plot the transition matrix among bout types [34] for the two states with the highest positive or negative log-fold change. The size of each edge represents the probability of visiting a state (we show only states with a probability *>* 0.1), and arrow widths are proportional to the transition probabilities (only transitions with probability *>* 0.1 are displayed). We find that *q*_*i*_ = 8 is enhanced in the naive fish, consisting of sequences of routine turns, high angle turns and shadow avoidance turns [34]. In contrast, fish previously raised with paramecia exhibit an enhancement of *q*_*i*_ = 10, which consists of sequences of Approach Swims and J-turns, elements of the canonical prey capture sequence [36]. (e) Complementary Cumulative Distribution Function (CCDF) of the time spent in *q*_*i*_ = 10 for the two conditions. We find an overall increase in bout-sequence duration following prior exposure to prey: the geometric mean duration was *τ*_N_ = 1.5 (1.45, 1.55) bouts for naive fish and *τ*_RW_ = 1.64 (1.58, 1.69) bouts for fish raised with paramecia, where values in parentheses indicate the 95% confidence interval obtained by bootstrapping across fish. The difference between conditions was significant in a two-sided permutation test p-value *<* 0.0005. The CCDF was obtained using all the available bout sequences, and error bars represent 95% confidence intervals bootstrapped across fish.

Focusing on the phenotypic variation across timescales, we find significant differences up till *q ≈*20 states, Fig. 6(b). To interpret the finest dynamical structure still supported by the data, we compare state usage in the *q* = 20 representation, Fig. 6(c),S6(g).

We find that naive fish more frequently visit states enriched in sequences of routine turns, high angle turns and shadow avoidance turns, whereas fish raised with paramecia increase their time spent in state *q*_*i*_ = 10, which is enriched in sequences of Approach Swims and J-turns, the canonical prey capture sequence [34, 36], Fig. 6(d). Not only do fish raised with prey visit state *q*_*i*_ = 10 more often, but they also increase the sequence length in this state, Fig. 6(e). Consistently, we find that fish raised with paramecia increase the time spent with eyes converged and reduce the time spent spent without eyes converged when compared to “naive” fish, Fig. S6(e1).

Exposure to rotifers showcases a different trend, Fig. S6(c,d), with fished raised with rotifers showing an enhancement of the time spent performing exploratory behaviors (sequences of slow forward movements and routine turns), and a reduction of the visits to *q*_*i*_ = 10, when compared to naive fish. They also spend more time without converged eyes, Fig. S6(e2), in contrast to the pattern observed when fish were raised with paramecia. We also find that the duration of sequences in state *q*_*i*_ = 10 is reduced for fish raised with rotifers, Fig. S6(f). Compared to previous analysis [16], our new approach uncovers a reduction of inter-individual differences upon prey exposure with paramecia, for which experienced fish deploy more canonical prey capture sequences.

## DISCUSSION

This study set out to faithfully reconstruct a space of variation among dynamical phenotypes in living systems. Comparing dynamical phenotypes requires deciding which differences in time matter, yet the relevant temporal resolution is generally unknown and constrained by finite observation time. We address this problem by defining a comparison across a multiscale transfer-operator descriptions, retaining dynamical differences only to the extent that they remain distinguishable from finite-sampling uncertainty. The resulting dissimilarity therefore lets the data determine the scales at which two dynamical realizations meaningfully differ. Because this dissimilarity defines a local metric on the underlying parameter space, its global diffusion-map coordinates can be interpreted in terms of directions of dynamical sensitivity rather than merely as axes of dimensionality reduction. We illustrated the approach in a toy model and then built phenotypic spaces for bacterial and larval zebrafish navigation, confirming known features of phenotypic variation in both systems but also uncovering previously unknown effects occurring at multiple time scales.

In bacteria, we used cell-position measurements from [27] to reconstruct a phenotypic space comprising isogenic wild-type cells together with cells in which CheR and CheB expression was experimentally varied. Phenotypic variation aligns much more strongly with CheB abundance than with CheR, consistent with the greater sensitivity to CheB reported previously [27]. This asymmetry cannot be explained simply by the antagonistic methylation and demethylation activities of CheR and CheB: Dufour et al. showed that CheB-dependent receptor deamidation is required to reproduce the enhanced sensitivity of tumble bias to CheB abundance, while slow fluctuations in receptor methylation further contribute to CheB-dependent behavioral variability. We further find that variation in run speed constitutes a major axis distinguishing individuals, consistent with previous work identifying run speed as an important component of non-genetic behavioral diversity [33]. Unexpectedly, run speed itself is strongly associated with CheB abundance, a relationship not directly predicted by the canonical chemotaxis network and whose mechanistic origin remains unclear. Wild-type and mutant cells also occupy different regions of the phenotypic space, with distinct behavioral strategies at its extremes. Cells with experimentally varied CheR and CheB levels are enriched near phenotypes that, within our search model, perform particularly well at either short or long spatial scales. Wild-type cells are instead enriched in a region characterized by more frequent fast runs and comparatively high search performance across a broad range of scales. This raises the possibility that the native regulation of CheR and CheB helps constrain phenotypic variation away from strategies specialized to particular environmental scales, which could be advantageous when environmental length scales fluctuate unpredictably [5]. Such an interpretation is consistent with the organization of the chemotaxis pathway: CheR and CheB are co-transcribed and translationally coupled, a regulatory architecture that buffers their relative expression and promotes robust chemotactic output [39, 40]. We emphasize, however, that the associations identified here, particularly between CheB abundance and run speed, remain correlative, and establishing their causal basis will require targeted perturbations.

In larval zebrafish, we analyzed bout-sequence data from [34]. Our previous analysis of this dataset was restricted to a single coarse-graining scale [16]. Here, by capturing coarse- and fine-scale differences within a single construction, we can resolve additional effects of prey experience on behavioral phenotypes. We found that prior exposure to paramecia substantially reduces the spread of individuals in phenotypic space. Thus, experience appears to affect not only the mean behavioral phenotype but also its variability: naive fish occupy a broader range of behavioral phenotypes, whereas prey-experienced fish converge toward a more stereotyped repertoire with bout sequences that support successful hunting. These fine-scale changes align with previous studies of hunting ontogeny: during prey capture, fish experienced to prey modulate their first turn towards the prey and the final capture maneuver to increase capture success [37, 38]. Our approach recovers these fine effects in bout sequences without relying on eye convergence measurements. At coarser scales, it also allows us to compare individuals over the full recording period, revealing that fish raised with paramecia spend less time performing exploratory behaviors.

Strikingly, these effects are prey-specific. In an assay with rotifers, prey-experienced fish instead spend more time performing exploratory behaviors, Fig. S6, while spending less time in the prey-capture state that is enhanced after rearing with paramecia. This observation is consistent with previous reports that fish raised with rotifers spend less time hunting while nevertheless hunting more efficiently than naive fish [38]. One possible explanation is that differences in prey motility allow rotifers to be captured more rapidly, leaving more time for exploratory behavior. Differences in the nutritional value of the two prey types could also contribute to this effect. Establishing whether larvae acquire genuinely prey-specific behavioral repertoires, and identifying the mechanisms responsible for such differences, will require additional replicates and controlled manipulations of prey exposure.

Across both systems we encounter substantial structure in phenotypic variation, with implications beyond the specific organisms studied. Despite the large number of underlying parameters that can in principle vary — protein expression levels in bacteria, or membrane properties, synaptic weights and neuromodulatory states in zebrafish — the observed behavioral variation is concentrated along only a few dominant dimensions. Previous work has shown that evolutionary constraints can restrict phenotypes to low-dimensional manifolds [22, 23], suggesting that population-level variation may reveal the effective directions of parameter variation that remain accessible. Furthermore, in both bacteria and zebrafish, we find that individuals are organized within simplex-like spaces whose extremes correspond to distinct behavioral strategies. Such geometries are reminiscent of Pareto-front models, in which trade-offs among competing objectives organize phenotypes between a small number of archetypal strategies [24, 25].

Methodologically, our approach builds on a rich history of reconstructing the intrinsic geometry of data points in manifold learning [41–44], and on work dissecting the manifold structure of phenotypic spaces in evolutionary contexts [6, 23]. Our methodological contribution is twofold: a new multiscale dissimilarity between nonlinear dynamical systems, and a connection between the diffusion map eigenvectors and the dominant modes of the metric tensor mapping parameter variation to differences in emergent dynamics. The latter connects our work to the notion of “sloppiness” in model inference [45], in which parameter combinations that most strongly affect model predictions are termed “stiff”; in the SI we show that these locally defined stiff modes guide the macroscopic modes of variation recovered by diffusion maps.

Recent approaches compare dynamics either by learning invariant trajectory representations or by asking whether two systems are dynamically equivalent. Contrastive Cartography, for example, uses contrastive learning to construct coordinate-invariant representations of stochastic trajectories and, when applied to the same *E. coli* dataset, recovered variation associated with tumble bias and CheB/CheR expression [46]. Dynamical similarity methods instead quantify how far systems are from being topologically conjugate [47–50]; our Perron-Frobenius operator approximation [13] (adjoint to Koopman [51]) also provides an efficient route to such comparisons, as we illustrate for the Lorenz system in the SI. Our approach addresses a complementary question: it explicitly compares coarse-grained dynamics across resolutions, calibrating each difference against finite-sampling uncertainty while retaining the identity of metastable states. This is important for behavioral variability, where individuals may visit the same states but differ in how strongly particular states are occupied [8, 52].

Three features of our construction bound its applicability. First, estimating transfer operators assumes that the dynamics are approximately stationary over the observation window, which is often not the case [15]. We thus essentially take an adiabatic approximation of the dynamics; longer recordings may be better described as trajectories through the phenotypic space, which would in turn allow within-individual change to be studied with the same machinery. Second, our multiscale dissimilarity requires a family of coarse-grained descriptions of the dynamics, and we obtain these with a deliberately simple and agnostic procedure based on clustering the eigenvectors of the reversibilized transfer operator and conditioning the solution on the forward coherence of each metastable state (see Methods). This was sufficient for all systems studied here, and the uncertainty scale against which we assess significant differences further regularizes against small errors in the coarse-graining. Other schemes are nonetheless available: Generalized Perron Cluster-Cluster Analysis [53], for example, is constructed to handle non-reversible dynamics with cyclic structures and may be better suited to certain classes of systems. Third, the construction assumes that individuals share a common state space, so that their coarse-grained operators can be directly compared. When realizations occupy genuinely different state spaces, as in comparisons across species, dynamical similarity analysis may be more appropriate [49, 54, 55], and our operator estimation provides an efficient route for this, as mentioned above.

Beyond these methodological limitations, the multi-scale structure could also be exploited for objectives other than the unsupervised reconstruction considered here. In the present work, differences across resolutions are weighted by their statistical resolvability, without using experimental labels. When predefined conditions are available, one could instead take a supervised approach and assess which dynamical resolutions are most informative for distinguishing among conditions.

A further caveat concerns the interpretation of the space itself. We have described the variation we recover as inter-individual, but in the datasets we studied each individual contributes a single finite-time observation, so what we strictly measure is variation among realizations. Establishing that this reflects consistent individual differences, requires repeated measurements of the same individual, the standard route in behavioral ecology [7, 52, 56]. Our significance testing ensures that the differences we retain exceed what finite sampling of identical dynamics would produce, which rules out the trivial case, but it does not establish that an individual would return to the same position in the space if recorded again. Nonetheless, our framework is well suited to these questions given the appropriate data. Repeated recordings would place each individual at several points in the space, allowing within-individual spread to be compared directly against between-individual spread. Following the same individuals through development would reveal whether animals occupy fixed positions, converge as they gain experience, or move along the manifold in a stereotyped way, and recording the same animals under different prey or environmental conditions would show whether the axes we recover are preserved across contexts or are specific to each assay.

The phenotypic spaces we reconstruct are a starting point to further investigate complex mechanisms of individual variability or evolution. Because phenotypic spaces are built directly from observed dynamics, they can be aligned with independent measurements of the parameters presumed to generate variation: single-cell expression measurements in bacteria, or whole-brain imaging in larval zebrafish, where experience-dependent improvements in hunting have already been traced to identified forebrain circuits [37]. An individual’s position in the space could then be mapped onto its internal state directly rather than inferred, and ultimately used to predict how it will respond to a perturbation rather than to describe variation after the fact. Combined with long-term evolution experiments [57] and comparisons across strains and species [8], this offers a route to studying evolutionarily driven behavioral variation with increased precision. In each of these settings, the variability that averaging discards is structured, and this structure is informative about the constraints shaping the parameters that control behavior.

## METHODS

### Data-driven coarse-graining of the dynamics

To infer coarse-grained Markov models from time series data, we start by approximating the Perron-Frobenius operator using the framework of maximally predictive Markov models [13]. When working from toy models, we directly use the full state space. When working from real data, we first reconstruct the state space using a time-delay embedding with *K* delays. We then partition the state space into *N* microstates *s*_*i*_,*i ∈* [1, 2, …, *N* ] using a k-means clustering. We set *K* by minimizing the unpredictability of the Markov chain (as measured through its entropy rate), and *N* as the largest partition size before finite-size effects in the estimation of the entropy rate become evident [13]. We then build a Markov transition matrix by counting transitions between microstates, *P*_*ij*_(*τ* ) = *P* (*s*_*j*_(*t*+*τ* ) *s*_*i*_(*t*)), yielding a discrete approximation of the Perron-Frobenius operator. We set the transition time *τ* such that the implied timescales *t*_imp_(*τ* ) of the inferred Markov chain are nearly invariant to *τ*, signaling Markovianity [13, 14, 16]. Using the fine scale transition matrices, we then proceed to identify coarse-grained metastable states. To use a common coarse-graining approach across all dynamical systems studied here, we use a simplified method that simultaneously takes advantage of the real eigen-spectrum of reversibilized transition matrices, while also optimizing for the metastability of the forward dynamics. Combining ideas from [18, 19, 28], we rescale the eigen-vectors of the reversibilized transition matrix by their respective eigenvalue, and then cluster this space into *q* states using a k-means clustering weighted by the invariant measure. While we work with the real eigenspectrum of reversibilized operators, we then try to find a coarse-graining the respects the metastability of the forward dynamics by repeating the clustering from random initial conditions (1000 times), and saving the *q* states that maximize the minimum metastability among all states. We assess metastability by estimating how much of the probability density remains in a state *q* after a timescale *τ*,

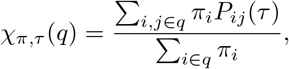

where *π* is the invariant density, *Pπ* = π.

Below we provide details of the transfer operator reconstruction for each dataset used in the main text.

#### Overdamped triple-well system

To construct an ensemble transition matrix, we combine data from all parameter values. Since the dynamics is memoryless, we skip the time-delay embedding step and directly partition the measurement time series into *N* = 250 for each experimental timescale *T*_expt_. We then set *π* = 0.05 s according to Fig. S1(a). In Fig. S1(b,c), we show the first two eigenvectors of the reversibilized transition matrix, and the obtained coarse-graining into *q* = 2 and *q* = 3 states that recover the wells of the system.

#### Bacterial motility

To compensate for the uneven sampling of different conditions, we construct ensemble transition matrices by sampling 10^4^ s from both wildtype and mutant cells, and then averaging the subsequent transition matrices for each *K* and *N* across 100 random seeds. Using the entropy rate of the Markov chain, we find that *K*^*\**^ = 0.5 s maximizes the predictability over the future, and *N*^*\**^ = 1700 is the largest number of partitions before finite-size effects, Fig. S3(a2). We choose *τ*^*\**^ = 0.2 s to respect the Chapman-Kolmogorov condition, Fig. S3(a3).

#### Zebrafish behavior

To compensate for the uneven sampling of different sensory contexts, we proceed as in [16] and to construct ensemble transition matrices by randomly sampling 7500 bouts from each condition for 100 seeds and then averaging the subsequent transition matrices. We use the same parameters as in [16], namely *K*^*\**^ = 5 bouts bouts, *N*^*\**^ = 1200 and *τ*^*\**^ = 3 bouts.

### Estimating the uncertainty scale for each distance measurement

We utilize a permutation test to determine if two realizations *α, β* were generated from the same underlying Markov chain at every *q*. From each pair of *q*-state symbolic sequence, *α* and *β*, we collect sequences of states separated by a time *τ*. From this pool of transition events, we then randomly sample the same number of transitions as observed in each realization *α* And *β*. Through this procedure, we obtain resampled Markov chains, 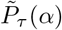 and 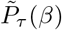, that respect the null hypothesis that the symbolic sequences in *α* and *β* were generated by the same Markov model. We then estimate the average distance between resampled Markov chains using Eq. 1, repeating this process for 100 random samples and averaging across all instances, Eq. 2. To avoid numerical division issues, if any *ε*_*q*_(*α, β*) ≈0, we regularize it to the maximum between the 5th percentile of the distribution of resampled distances or 10^−5^.

### Self-tuning diffusion maps

To create a self-tuning diffusion map, we follow the methodology from [58]. After the estimation of the multiscale dissimilarity, Eq. 3, *D*_*α, β*_ = *D*(*α, β*), we transform it into a heat kernel,

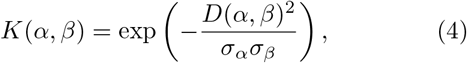

with a self-tuning bandwidth *σ*_*α*_ *σ*_*β*_ [58]. This band-width is the distance to *k*-th nearest neighbor, 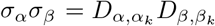, where *α*_*k*_ is *k*-th nontrivial nearest neighbor for point *α*. This approach is robust to outliers while preserving local structure [58].

To remove the effect of the data distribution and retrieve an anisotropic kernel, we transform the above expression by dividing by the empirical estimate of the density,

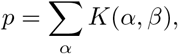

getting the new collumn-normalized kernel,

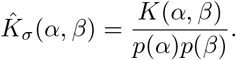

We then row-normalize this kernel to find the anisotropic transition kernel *T* (*α, β*). The two normalization steps ensure that the embedding is insensitive to how densely phenotypes are sampled, so that the recovered geometry reflects the structure of the phenotypic manifold rather than the distribution of individuals on it [26].

We then obtain the diffusion coordinates by computing the non-trivial eigenvectors of *T* (*α, β*), *ψ*_*m>*0_, normalizing them by the stationary distribution and rescaling by their respective eigenvalues *λ*_*m>*0_, **v**_*m*_ = *λ*_*m*_*ψ*_*m*_. These diffusion coordinates define the effective axes or variation and are subsequently used for visualizing and clustering the phenotypic space. *ψ*

To select the appropriate *k*-th nearest neighbor we rely on inspecting the eigenvalue spectrum of *T* (*α, β*) as *k* increases. In Fig. S2(d1-d4), we showcase how in the triple well system of Fig. 2 choosing different *k* captures different properties of the data. A low *k* = 20, the diffusion map has 3 non-trivial eigenvalues with eigenvectors that cleanly separate out the 4 clusters in the data. Such a representation is useful for spectral clustering. As *k* increases, the global structure begins to emerge with 2 dominant non-trivial eigenvalues. In Fig. S7, we report the eigenvalue spectra for various *k* for both the bacteria, Fig. S7(a) and larval zebrafish, Fig. S7(b), datasets. The error bars represent 95% confidence intervals that were estimated by resampling 50% of the data over a 100 random seeds. We set *k* when the eigenvalue separation is sufficiently strong, which corresponds to approximately 10% of the number of data points in each dataset.

### Discovering the phenotypic groups using fuzzy c-means

To select out individuals in each extreme of the phenotypic space, we perform weighted fuzzy c-means clustering on the first two diffusion coordinates (**v**_1_, **v**_2_). The fuzzy c-means is weighted by density of the data points, approximated by the eigenvector of eigenvalue 1 of the diffusion operator.

### Symbolic sequence and trajectory simulations for phenotypic groups

We utilize the predictive power of our Markov transition matrices to simulate artificial trajectories for both the bacterial and larval zebrafish behavior. We begin by calculating ensemble Markov transition matrices for each phenotypic group 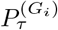 sutilizing only the data points belonging to group *G*_*i*_. In order to retain all the finescale information, we estimate 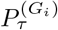 at the shortest τ possible for each setting. For bacterial navigation this corresponds to τ = 0.01*s* and for larval zebrafish this corresponds to τ = 1 bout. We then simulate long sequences of microstates *s*_*i*_,*i* ∈ 1, …, *N*^*^ according to this transition matrix 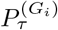, sampled from a random initial microstate. For bacteria, this corresponds to 10000 sequences 500 s long at a sampling rate of 10Hz per phenotypic group. For larval zebrafish, this corresponds to 10000 sequences 5000 bouts per phenotypic group. From each simulated sequence of microstates, we generate lab space trajectories by sampling velocity vectors from the real-data distribution of vectors within each microstate.

### Resource detection estimation from simulated trajectories

Using the simulated lab space trajectories, we estimate and compare the efficiency of detecting resources. For the larval zebrafish, we perform the estimation in two different conditions from [16]. In the first condition, we assume that the fish is converging its eyes and executing the prey pursuit sequence. Thus, the task is to collect resources uniformly distributed in a 60^°^ aperture [35] at a maximum distance of *r* = 2mm [36] from the fish. We also consider that the fish captures prey only if its head position is within a short distance of *l*_*hunt*_ = 0.05 body lengths. In the second condition, we assume that the fish are detecting resources that are distributed densely and uniformly in a radius of *r* around them. We assume that fish can detect resources (visually, acoustically, or mechanically) within a distance of *l*_detect_ = 1 body length from them.

For the bacteria, we only estimate the condition of detecting resources that are distributed densely and uniformly around them in a radius *r*. While bacteria do not perform spatial sensing, we set *l*_detect_ = 10*µm*, which roughly corresponds to the size of the bacteria plus their flagellum. We perform this calculation solely to assess the chance that the bacteria will detect changes in a fictitious chemical gradient at a distance *r* from it.

### Assessing variability through phenotypic space entropies

To assess whether raising fish with prey reduces inter-individual differences, we estimated the entropy of the distribution of fish along the dominant modes (**v**_1_, **v**_2_). To do that, we approximated the probability distributions using a 2D Gaussian kernel density estimate, with a bandwidth of 0.1, which is approximately twice the median nearest neighbor distance among fish hunting paramecia. Since each condition had a different number of fish, we resampled without replacement from the condition with most fish to obtain an equal-*n* sampling that matches the number of fish in the smaller condition. We then obtained 95% confidence intervals by bootstrapping among equal size samples. Significance estimates were obtained using a one-sided permutation test on equal size samples. We found that for fish hunting paramecia (*n* = 148 fish across the two conditions), the significance results are insensitive to the choice of bandwidth (varying the bandwidth between 0.05 and 0.3 always yields a p-value *<* 0.001). However, for fish hunting rotifers (*n* = 46 fish across the two conditions), the difference in entropies is not consistently significant across band-widths, possibly reflecting the small sample size.

### Datasets and toy models

#### Overdamped Langevin dynamics from a 3 well potential

We simulate time series from an overdamped Langevin equation of the following form,

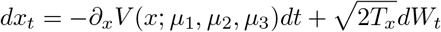

where *V* (*x*; *µ*_1_, *µ*_2_, *µ*_3_) is the potential driving the dynamics, *T*_*x*_ is the thermal noise level and *W*_*t*_ is a Wiener process. For the phenotypes, we generate 4 unique potentials by fitting a spline using a Piecewise Cubic Hermite Interpolating Polynomial (PCHIP) to a 3 variable local minima *θ*_*i*_ = *V* (*µ*_*i*_) and 4 fixed local maxima. We set *θ*_3_ as a function of *θ*_2_, restricting variation to a plane. The height of the minima is sampled from a gaussian distribution with a standard deviation of 0.01 and varying means. For the first minimum, *V* (*µ*_1_), the mean is -2 for the blue and orange phenotypes and -0.2 for the green and red phenotypes. For the second minimum, *V* (*µ*_2_), the mean is -1.2 for the blue and red phenotypes and -1 for the green and orange phenotypes. The height of the third minimum is set as *V* (*µ*_3_) = −0.98*V* (*µ*_2_) −2.22 for each of the phenotypes. In this way, blue and orange, and red and green share *V* (*µ*_1_), whereas red and blue, and orange and green share *V* (*µ*_2_) and *V* (*µ*_3_). We then generate a variable number of realizations from each of the four phenotypes (25 realizations for the blue phenotype, 29 for the orange, 24 realizations for the green and 28 realizations for the red) with initial conditions drawn uniformly between -1 and 1 with *T*_*x*_ = 0.5, *δt* = 0.05*s* for a total time *T*_expt_ = 10^4^s. For Fig. S2, we sample simulation durations from 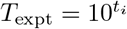in the range *t*_*i*_ ∈ [1, 4].

#### Bacterial (Escherichia coli) navigation dataset

We utilize the dataset from [27] where the locomotion of isogenic wild-type bacteria and mutants is recorded. In this dataset, wild-type RP437 *E. coli* cells make up the wild-type dataset and are also the parental strain for all the mutants with fluorescent protein expression. The fluorescent cells are made by first genetically constructing fluorescent protein fusions CheB-mYFP and mCherry-CheR. The gene constructs are then placed under the control of two independent inducible promotors (Rhamnose or Isopropyl *β*-D-1-thiogalactopyranoside) on the chromosome in a strain lacking both the native cheR and cheB genes (see [27] for details). Cells are grown to mid-exponential phase in minimal medium and then suspended in motility buffer, in which the behavior remains nearly constant for more than one hour [27, 59]. The cells are kept under a 22 mm^2^ cover slip with ≈10*µm* in depth, confining their motion to quasi-2D. Cell motility was recorded at 10Hz temporal resolution in a 1.3mm square field of view, with a spatial resolution of 169 *µm*/pixel. The tracks are then extracted using custom MATLAB code [27]. The recorded (*x, y*) positions of the cells were used to calculate velocity auto-correlation *C* _ν_. Specifically, the velocity autocorrelation is fit using a nonlinear least squares method to the functional form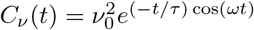, where *ν*_0_ is the average cell speed, τ is the cell directional persistence and ωis the angular frequency of the circular trajectory of the cell. If ω is particularly high, it is indicative of the cells swimming close to the coverslip surface. For our analysis, we consider only tracks that are longer 10 s, and with a fit 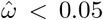 rad. For the fluorescent cells dataset, we noticed the cells appeared immobile for the last 10 s of most recordings. We thus clipped the last 10 s of all recordings for the fluorescent cells. We also subsequently noticed that in many cells the interaction with the cover slip was too brief to be detected by the autocorrelation criterion, but long enough for our approach to be sensitive to it. As this circling behavior does not seem indicative of naturalistic locomotion, we subsequently removed 83 tracks from the RP437 dataset and 48 tracks from the fluorescent cells dataset upon manual inspection. Finally, in total we used 5399 RP437 tracks and 1374 fluorescent cell tracks. The protein expression of the fluorescent cells was measured using single-cell fluorescence microscopy [27]. The behavior of each cell is represented by its egocentric velocities *v*_||_ and *v*_⊥_ as introduced in [60]. In brief, we use 3 points in the trajectory along time points *t*_−1_, *t*_0_, *t*_1_. We project the vector from *t*_0_ to *t*_1_ onto the vector along *t*_*−*1_, *t*_0_. Thus, *v*_||_ represents forward of backward motion and *v*_⊥_ represents left or right motion with respect to the motion at the previous time point. These features are then standardized across the entire dataset to have zero mean and standard deviation of 1.

#### Zebrafish (Danio rerio) navigation dataset

We leverage the dataset from [34] and previously analyzed in [16]. Wild type Tubingen larvae of ages 6 − 7 days post fertilization were recorded at high temporal resolution (700Hz) at two different pixel sizes - 27*µm*/pixel for exposed to prey conditions and 58*µm*/pixel for not exposed to prey conditions. 8 points on the tail of the fish are tracked during the experiment using a custom online tracking software [34]. Same as [16], we utilize 463 fish from 14 different sensory contexts, retaining only those detected bouts that generated a velocity of at least 4mm/s. Each bout is a 250 ms ×8 dimensional object, which we project onto 20 dimensions using principle component analysis as in [16]. We then work with the sequences of bouts to reconstruct the state space. We also utilize the eye convergence index (bout with or without eye convergence) for each bout recorded for the fish exposed to prey as a marker of hunting attempts, which is described in detail in [16, 34]. The details of each of the sensory context is provided below, from [16]:

**Light 5**× **5** cm^**2**^: Fish were presented with a uniform light from below with an illuminance of 1000 lm/m^2^ in a 5 ×5 cm^2^ squared arena with 3 mm of depth.

**Light 1**× **5** cm^**2**^: Fish were presented with a uniform light from below with an illuminance of 1000 lm/m^2^ in a 1 × 5 cm^2^ squared arena with 8 mm of depth.

**Dark 5** × **5** cm^**2**^: Fish were presented with darkness (0 lm/m^2^) in a 5 ×5 cm^2^ squared arena with 3 mm of depth.

**Expanding Spot 5** ×**5** cm^**2**^: An expanding dark spot at different speeds (0.25, 0.5, 1, 1.5, 2.0, 2.5 cm/s) and different orientations (0^*°*^, 90^*°*^, 180^*°*^, 270^*°*^) was presented in closed loop 4 cm away from the larva. Stimuli were randomized and presented every 2 min. This assay was done in a 5 ×5 cm^2^squared arena with 0.3 cm of depth.

**Dark Transitions 5** ×**5** cm^**2**^: The spontaneous swimming with light transitions assay was based on [61]. Fish were left in the dark for 30 minutes and then presented with uniform light at different intensities (0, 12, 44, 104, 232, 447, 790, 1890, 4700 lm/m^2^) for 3 min in a 5× 5 cm^2^ arena with 3 mm of depth.

**Phototaxis 5**× **5** cm^**2**^: The phototaxis assay is based on [62] and performed in 5× 5 cm^2^ arena with 3 mm of depth. The stimulus consists of a uniform brightness of varying intensity (100, 410, 780, 1250 lm/m^2^) on one side of the fish and darkness on the other. The stimulus was in closed loop with the larva for a duration of 12 s.

**Forward Optomotor response 1**× **5** cm^**2**^: The forward optomotor response assay was performed as described in [63], but using a 1 ×5 cm^2^ with 8 mm depth arena. Drifting gratings with a spatial period of 1 cm and ten different speeds (0, 2.5, 5, 7.5, 10, 15, 20, 30, 40, 50 mm/s) were presented from below when the larva is at the extremities of the arena. Trials end when fish reach the opposite end of the arena. After, there is 5s of intertrial interval of homogenous light (1000 lm/m^2^) and a new trial starts with gratings in the opposite direction. In trials that larvae are not able to reach the opposite end of the arena (*>* 30 s) a 10 mm/s grating was displayed until it swam the remaining distance.

**Directional Optomotor response 5**× **5** cm^**2**^: The directional optomotor response assay was performed as in [64]. A **5**× **5** cm^**2**^ arena with 3 mm of depth was used. Drifting gratings with a spatial period of 1 cm, moving at 10 mm/s, and from 24 different orientations (15° apart) were presented from below to the larva and in closed loop with its orientation. Trials last 10s and start when the larva wass in the center of the arena. During the inter-trial interval, that would last at least 5s, circular converging gratings were projected to drive the larva to the center of the arena.

**High Lux light-dark transitions 5 × 5** cm**2**: Fish were exposed to alternating 3 min periods of high illuminance light (5000 lm/m2) and darkness in an 5 ×5 cm2 arena with 3 mm of depth.

**Prey Capture 2.5× 2.5** cm**2**: All prey capture datasets were performed in 2.5 ×2.5 cm2 arenas with 3 mm depth and larvae were illuminated from above (1000 lm/m2). Fish were fed with 50-100 paramecia (*Paramecium caudatum*) or rotifers (*Brachionus plicatilis*) and were allowed to hunt for 1-2h. A subset of fish were never fed until the assay, while others are raised, starting at 3 days post fertilization, with the type of prey that is used in the assay. To ensure that these fish were not satiated they are starved for at least 2 hours before the assay.

**Freely exploring in the light, raised with rotifers 2.5** ×**2.5** cm**2**: Fish were fed with 50-100 rotifers starting at 3 days post fertilization and then placed in 2.5 cm2 arenas with 3 mm depth and illuminated from above (1000 lm/m2) to freely swim. To ensure that these fish were not satiated they were starved for at least 2 hours before the assay.

## CODE AND DATA AVAILABILITY

The code used to generate the results and figures in this work is freely available on Github: https://github.com/GautamSridhar/Multiscale_phenotypic_variation. The data and the analysis results are freely available on Kaggle: https://doi.org/10.34740/kaggle/dsv/20089164

## ACKNOWLEDGEMENTS

We would like to thank the Wyart lab “SIBBIL” for their inputs. We would also like to thank Thierry

Emonet, David Jordan and João Costa, for constructive discussions that helped shape the manuscript. We thank the 2020 Fondation Bettencourt-Schueller (FBS-don-0031) award (“Identity and organization or neuronal networks controlling exploration”), the 2020 Prize Equipe “Fondation pour la Recherche Médicale” (FRM-EQU202003010612) ‘Neuronal circuits underlying navigation: from genes to behavioral models’, the 2020 European Research Council Consolidator grant no. 101002870 (2021–2026) “Exploratome: Circuit mechanisms underlying sensory-evoked navigation” awarded to CW., the European Union’s Horizon 2020 research and innovation program under a Marie Sk$lodowska-Curie grant no. 813457 ZENITH awarded to CW. that covered the salary of GS. The project benefited as well from the support of the Agence Nationale pour la Recherche (ANR) ANR-22-CE37-0023 LOCOCONNECT, ANR-23-CE16-0017-02 RocSMAP, ANR-24-CE16-7992 CIRCOLOCO, ANR-21-CE14-0042 MOTOMYO and ANR-21-CE13-0008 ASCENTS. GS was also partially supported by funds from OIST Graduate University. ACC acknowledges support from the Agence Nationale pour la Recherche (ANR) with the project ANR-25-CE02-1309. This research was supported in part by the Richard Mille Foundation and the National Science Foundation (NSF) grant PHY-2309135 to the Kavli Institute for Theoretical Physics (KITP) and the Gordon and Betty Moore Foundation Grant No. 2919.02.

## Author contributions

G.S., C.W. and A.C.C conceived the study. G.S. and A.C.C developed the methods and analysis code, prepared figures and performed mathematical modeling. G.S. and A.C.C. wrote the original draft. All authors reviewed and edited the manuscript. C.W. and A.C.C. supervised the research and secured funding.

## Ethics declaration

not applicable.

## References

[1] J. Golowasch, M. S. Goldman, L. Abbott, and E. Marder, Journal of neurophysiology 87, 1129 (2002).

[2] C. Pantoja, A. Hoagland, E. C. Carroll, V. Karalis,A. Conner, and E. Y. Isacoff, Neuron 91, 587 (2016).

[3] W. Löscher, Eneuro 11 (2024).

[4] A. Sih, A. Bell, and J. C. Johnson, Trends in ecology & evolution 19, 372 (2004).

[5] N. W. Frankel, W. Pontius, Y. S. Dufour, J. Long,L. Hernandez-Nunez, and T. Emonet, Elife 3, e03526 (2014).

[6] S. M. Huttegger and P. Mitteroecker, Evolutionary Biology 38, 335 (2011).

[7] K. L. Laskowski, D. Bierbach, J. W. Jolles, C. Doran, and M. Wolf, Nature Communications 13, 6419 (2022).

[8] D. G. Hernández, C. Rivera, J. Cande, B. Zhou, D. L. Stern, and G. J. Berman, Elife 10, e61806 (2021).

[9] B. de Bivort, S. Buchanan, K. Skutt-Kakaria, E. Gajda, J. Ayroles, C. O’Leary, P. Reimers, J. Akhund-Zade,R. Senft, R. Maloney, et al., Frontiers in Behavioral Neuroscience 16, 836626 (2022).

[10] S. R. Datta, D. J. Anderson, K. Branson, P. Perona, and A. Leifer, Neuron 104, 11 (2019).

[11] A. E. Brown and B. De Bivort, Nature Physics 14, 653 (2018).

[12] G. J. Berman, BMC biology 16, 1 (2018).

[13] A. C. Costa, T. Ahamed, D. Jordan, and G. J. Stephens, Chaos: An Interdisciplinary Journal of Nonlinear Science 33, 023136 (2023).

[14] A. C. Costa, T. Ahamed, D. Jordan, andG. J. Stephens, Proceedings of the National Academy of Sciences 121, e2318805121 (2024), https://www.pnas.org/doi/pdf/10.1073/pnas.2318805121.

[15] A. C. Costa, G. Sridhar, C. Wyart, and M. Vergassola, PRX Life 2, 023001 (2024).

[16] G. Sridhar, M. Vergassola, J. C. Marques, M. B. Orger,A. C. Costa, and C. Wyart, Proceedings of the National Academy of Sciences 121, e2410254121 (2024).

[17] E. M. Bollt and N. Santitissadeekorn, Applied and computational measurable dynamics (SIAM, 2013).

[18] G. Froyland and M. Dellnitz, SIAM Journal on Scientific Computing 24, 1839 (2003).

[19] G. Froyland, Physica D: Nonlinear Phenomena 200, 205 (2005).

[20] J.-H. Prinz, H. Wu, M. Sarich, B. Keller, M. Senne,M. Held, J. D. Chodera, C. Schütte, and F. Noé, The Journal of chemical physics 134 (2011).

[21] E. Marder, Eneuro 10 (2023).

[22] T. U. Sato and K. Kaneko, Physical Review Research 2, 013197 (2020).

[23] K. Kaneko, Journal of Statistical Mechanics: Theory and Experiment 2024, 024002 (2024).

[24] O. Shoval, H. Sheftel, G. Shinar, Y. Hart, O. Ramote,A. Mayo, E. Dekel, K. Kavanagh, and U. Alon, Science336, 1157 (2012).

[25] H. Sheftel, O. Shoval, A. Mayo, and U. Alon, Ecology and evolution 3, 1471 (2013).

[26] R. R. Coifman and S. Lafon, Applied and computational harmonic analysis 21, 5 (2006).

[27] Y. S. Dufour, S. Gillet, N. W. Frankel, D. B. Weibel, and T. Emonet, PLoS computational biology 12, e1005041 (2016).

[28] F. Noé and C. Clementi, Journal of chemical theory and computation 11, 5002 (2015).

[29] J. L. Spudich and D. E. Koshland, Nature 262, 467–471 (1976).

[30] H. Park, W. Pontius, C. C. Guet, J. F. Marko,T. Emonet, and P. Cluzel, Nature 468, 819–823 (2010).

[31] H. C. Berg, E. coli in Motion (Springer, 2004).

[32] G. H. Wadhams and J. P. Armitage, Nature Reviews Molecular Cell Biology 5, 1024–1037 (2004).

[33] M. Pleška, D. Jordan, Z. Frentz, B. Xue, and S. Leibler, Proceedings of the National Academy of Sciences 118, e2023322118 (2021).

[34] J. C. Marques, S. Lackner, R. Félix, and M. B. Orger, Current Biology 28, 181 (2018).

[35] I. H. Bianco, A. R. Kampff, and F. Engert, Frontiers in systems neuroscience 5, 101 (2011).

[36] M. B. McElligott and D. M. O’Malley, Brain Behavior and Evolution 66, 177 (2005).

[37] C. S. Oldfield, I. Grossrubatscher, M. Chavez,A. Hoagland, A. R. Huth, E. C. Carroll, A. Prendergast,T. Qu, J. L. Gallant, C. Wyart, et al., Elife 9, e56619 (2020).

[38] K. Lagogiannis, G. Diana, and M. P. Meyer, Elife 9, e55119 (2020).

[39] M. Kollmann, L. Løvdok, K. Bartholomé, J. Timmer, and V. Sourjik, Nature 438, 504 (2005).

[40] L. Løvdok, K. Bentele, N. Vladimirov, A. Müller, F. S. Pop, D. Lebiedz, M. Kollmann, and V. Sourjik, PLoS biology 7, e1000171 (2009).

[41] L. Van der Maaten and G. Hinton, Journal of machine learning research 9 (2008).

[42] S. T. Roweis and L. K. Saul, science 290, 2323 (2000).

[43] V. Roth, J. Laub, M. Kawanabe, and J. M. Buhmann, IEEE Transactions on Pattern Analysis and Machine Intelligence 25, 1540 (2003).

[44] L. McInnes, J. Healy, and J. Melville, arXiv preprint arXiv:1802.03426 (2018).

[45] M. K. Transtrum, B. B. Machta, K. S. Brown, B. C. Daniels, C. R. Myers, and J. P. Sethna, The Journal of chemical physics 143 (2015).

[46] N. Romeo, C. Chi, A. R. Dinner, and E. R. Jerison, PRX Life 3, 033022 (2025).

[47] M. Ostrow, A. Eisen, L. Kozachkov, and I. Fiete, Advances in Neural Information Processing Systems 36, 33824 (2023).

[48] A. Huang, M. Ostrow, S. H. Singh, L. Kozachkov, I. Fiete, and K. Rajan, arXiv preprint arXiv:2510.25943 (2025).

[49] S. Kamiya, J. Kitazono, and M. Oizumi, in ICLR 2024 Workshop on Representational Alignment (2024).

[50] S. Zhang, Z. Ye, Y. Yan, Z. Song, Y. Wu, and J. Wu, in Forty-second International Conference on Machine Learning (2025).

[51] J. N. Kutz, S. L. Brunton, B. W. Brunton, and J. L. Proctor, Dynamic mode decomposition: data-driven modeling of complex systems (SIAM, 2016).

[52] Z. Werkhoven, A. Bravin, K. Skutt-Kakaria, P. Reimers, L. F. Pallares, J. Ayroles, and B. L. De Bivort, Elife 10, e64988 (2021).

[53] B. Reuter, K. Fackeldey, and M. Weber, The Journal of Chemical Physics 150, 174103 (2019).

[54] A. E. Cohen, A. D. Hastewell, S. Pradhan, S. W. Flavell, and J. Dunkel, Physical Review Letters 130, 258402 (2023).

[55] T. Germain, R. Flamary, V. R. Kostic, and K. Lounici, arXiv preprint arXiv:2509.24920 (2025).

[56] K. L. Laskowski, C.-C. Chang, K. Sheehy, and J. Aguiñaga, Annual Review of Ecology, Evolution, and Systematics 53, 161 (2022).

[57] D. Chihoub, C. Pintard, R. E. Lenski, O. Tenaillon, and A. Couce, Proceedings of the National Academy of Sciences 122, e2501901122 (2025).

[58] L. Zelnik-Manor and P. Perona, Advances in neural information processing systems 17 (2004).

[59] V. Sourjik, A. Vaknin, T. S. Shimizu, and H. C. Berg, in Methods in enzymology, Vol. 423 (Elsevier, 2007) pp. 365–391.

[60] L. Tao, S. Ozarkar, J. M. Beck, and V. Bhandawat, Elife 8, e41235 (2019).

[61] H. A. Burgess and M. Granato, Journal of Experimental Biology 210, 2526 (2007).

[62] K.-H. Huang, M. B. Ahrens, T. W. Dunn, and F. Engert, Current Biology 23, 1566 (2013).

[63] K. E. Severi, R. Portugues, J. C. Marques, D. M. O’Malley, M. B. Orger, and F. Engert, Neuron 83, 692 (2014).

[64] M. B. Orger, A. R. Kampff, K. E. Severi, J. H. Bollmann, and F. Engert, Nature neuroscience 11, 327 (2008).

[65] I. Chavel, B. Randol, and J. Dodziuk, Eigenvalues in Riemannian Geometry, Pure and Applied Mathematics (Academic Press, 1984).

[66] P. Petersen, Riemannian geometry (Springer, 2006).

[67] E. N. Lorenz, Journal of Atmospheric Sciences 20, 130 (1963).

[68] J. Chang and G. Cooper, Journal of Computational Physics 6, 1 (1970).

[69] C. Sparrow, The Lorenz equations: bifurcations, chaos, and strange attractors (Springer Science & Business Media, 2012).

[70] P. Virtanen, R. Gommers, T. E. Oliphant, M. Haberland, T. Reddy, D. Cournapeau, E. Burovski, P. Peterson, W. Weckesser, J. Bright, S. J. van der Walt, M. Brett, J. Wilson, K. J. Millman, N. Mayorov, A. R. J. Nelson, E. Jones, R. Kern, E. Larson, C. J. Carey,İ. Polat, Y. Feng, E. W. Moore, J. VanderPlas, D. Laxalde,J. Perktold, R. Cimrman, I. Henriksen, E. A. Quintero,C. R. Harris, A. M. Archibald, A. H. Ribeiro, F. Pedregosa, P. van Mulbregt, and SciPy 1.0 Contributors, Nature Methods 17, 261 (2020).

[71] P. Frederickson, J. L. Kaplan, E. D. Yorke, and J. A. Yorke, Journal of differential equations 49, 185 (1983).

