## Supplementary Information for "Unraveling phenotypic variation in multiscale behavioral dynamics"

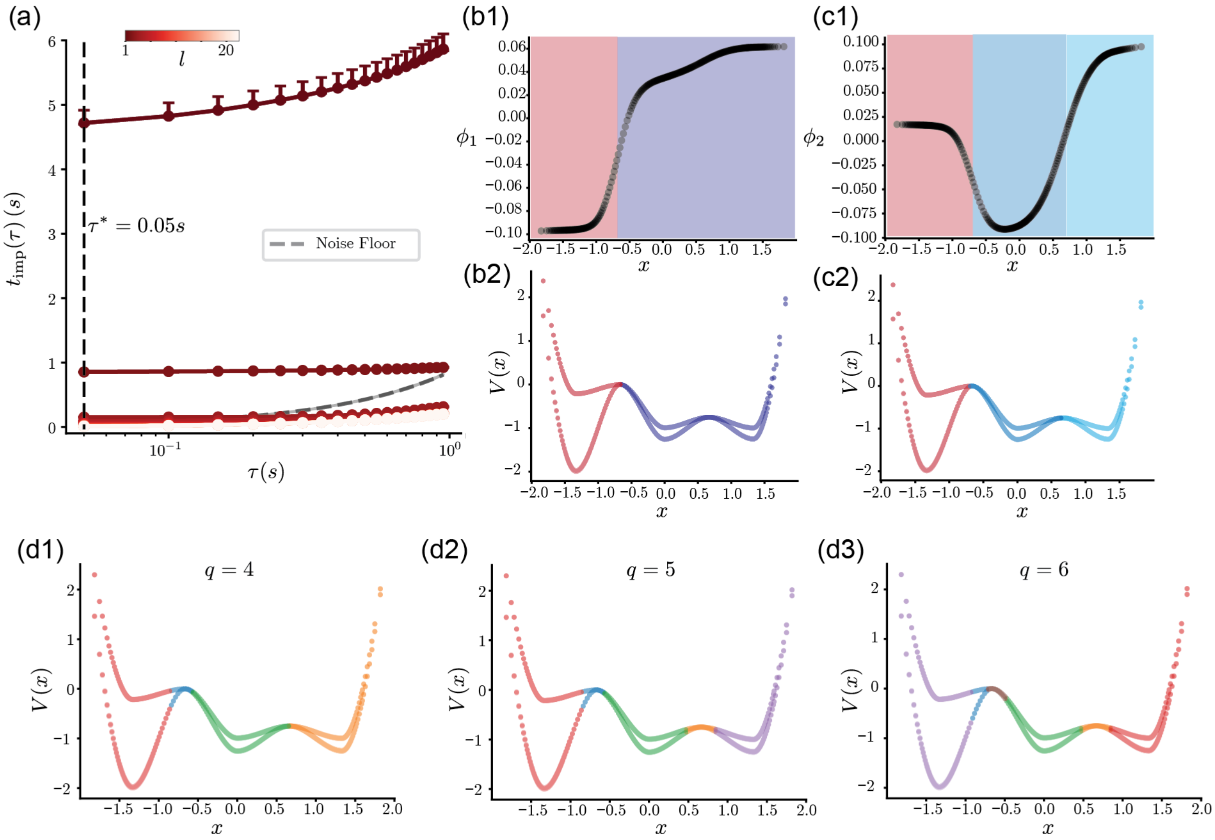


FIG. S1. **Details of the estimation of the ensemble transfer operator for the overdamped dynamics on a 3-well potential.** We first partition the state space consisting of all simulations across phenotypes into $N = 250$ microstates. (a) Implied timescales $t_{\mathrm{imp}}(\tau)$. We select $\tau = 0.05s$ to construct the ensemble transfer operator $P_{\mathrm{ensemble}}$ (b1) The first non-trivial eigenvector $\phi_1$ identifies the largest energy barrier in the system, enabling a coarse-graining into 2 states. (c1) The second non-trivial eigenvector $\phi_2$ further identifies the second-largest energy barrier in the system, resulting in a 3-state Markov chain capturing transitions among the metastable wells. (d1-d3) Coarse-graining for $q = 4, 5, 6$. Subsequent coarse-graining splits finescale dynamics in the transition regions while preserving the bulk of the metastable sets.

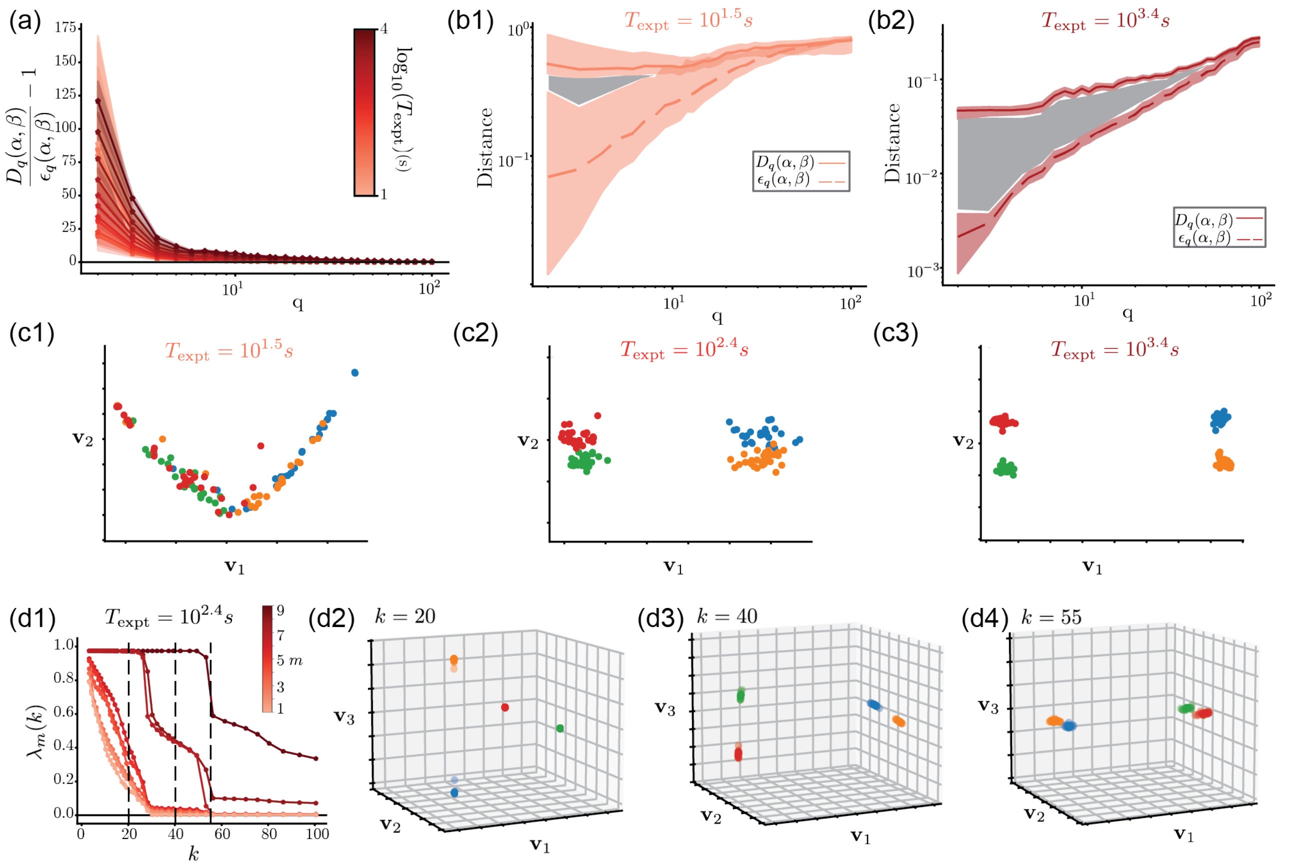


FIG. S2. **Dependence of the phenotypic space reconstruction on the simulation time $T_{\rm expt}$ and the number of nearest neighbors $k$, for the overdamped dynamics from a 3-well potential.** (a) We plot $D_q/\varepsilon_q - 1$ as a function of $q$, averaged across all observations. The area under each curve represents the multiscale distance metric $D(\alpha,\beta)$. Errorbars represent 95% confidence intervals across simulations. As $T_{\rm expt}$ increases, the area under the curve increases, pointing to increasing resolution in the differences between phenotypic groups. (b1,b2) Two examples of plots of $D_q(\alpha,\beta)$ and $\varepsilon_q(\alpha,\beta)$ at $T_{\rm expt} = 10^{1.5}$ s and $T_{\rm expt} = 10^{3.4}$ s. Shaded areas point to the multiscale distance that increases with $T_{\rm expt}$ (c) Projection along the first two diffusion coordinates $(\mathbf{v}_1, \mathbf{v}_2)$ (obtained with $k = 55$ nerest neighbors)) for different values of $T_{\rm expt}$. (c1) At low $T_{\rm expt} = 10^{1.5}$ s, the approach cannot distinguish blue from orange or red from green phenotypes, and yields an approximately 1D manifold. (c2) At $T_{\rm expt} = 10^{2.4}$ s, the first major axes are resolved, but there is still some overlap between phenotypes. (c3) At $T_{\rm expt} = 10^{3.4}$ s, all phenotypes are resolved completely. (d1-d4) Dependence of the diffusion map eigenvalues and coordinates on the number of nearest neighbors $k$ for $T_{\rm expt} = 10^{3.4}$ s. (d1) Eigenvalues $\lambda_m(k)$ as a function of $k$ for various eigenvalues $m$. We focus on 3 regimes - $k = 20$, for which the kernel bandwidth corresponds to within-cluster distances, $k = 40$, for which the kernel bandwidth now reaches the nearest cluster, and $k = 55$, for which the kernel bandwidth reaches all clusters. We notice a local vs. global information trade-off slowly unraveling as $k$ increases. (d2) For $k = 20$, the information is local, we receive $\lambda_{m\in 1,2,3} \approx 1$ and separated from the bulk. In this regime, the diffusion map eigenvectors $\mathbf{v}_1, \mathbf{v}_2, \mathbf{v}_3$ separate each cluster directly.(d3) For $k = 40$, the global access of variation begin to be resolved. (d4) At $k = 55$, the global information capturing 2 major axes of variation becomes clear.

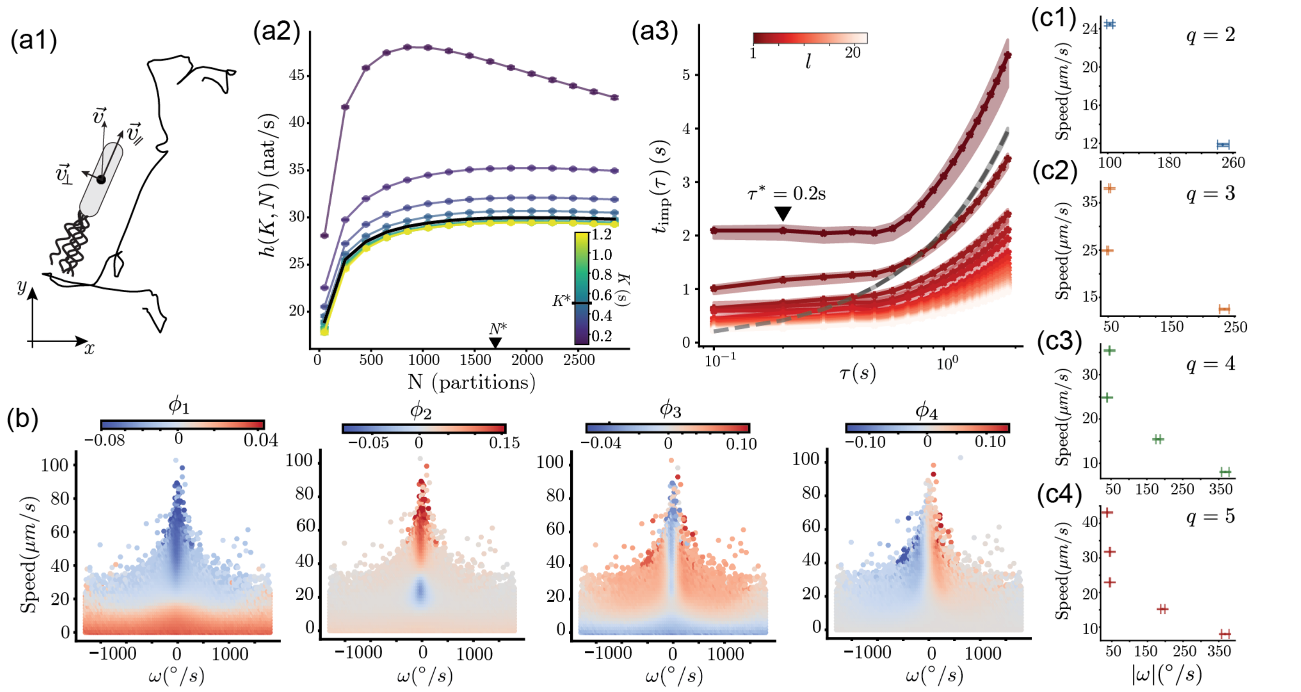


FIG. S3. **Details on the transfer operator approximation and coarse-grained dynamics for the bacterial chemotaxis dataset from [27].** (a1) We begin by estimating the parallel and perpendicular components of the velocity vectors estimated from the centroid trajectories. (a2) To ensure a maximally-predictive Markovian description, we perform a time delay embedding, and choose the number of delays $K$ by minimizing the entropy rate of the resulting Markov chain [13] (see Methods). We select $K^* = 0.5\,$s to minimize the entropy rate, and $N^* = 1700$ microstates to maximize the resolution over the dynamics before finite-size effects impair the estimation of the entropy rate. (a3) Implied timescales $t_{\mathrm{imp}}(\tau)$. We select $\tau^* = 0.2s$ to construct the ensemble transfer operator, in a regime where the inferred timescales are nearly invariant to $\tau$, indicating Markovianity [13]. (b) We report the first $\phi_{l\in[1,4]}$ non-trivial eigenvectors: $\phi_1$ differentiates running and tumbling behavior, $\phi_2$ differentiates an intermediate speed state from running and tumbling, $\phi_3$ differentiates smooth turning behaviors from straight runs and tumbles, and $\phi_4$ differentiates left-biased runs from right-biased runs. (c) Mean speeds and absolute changes in orientation $|\omega|$ for $q \in [2, 5]$. (c1) At $q = 2$, we identify running and tumbling states. (c2) At $q = 3$, the run states splits into fast and slow. (c3-c4) As $q$ increases, we find finer scale partitions of the running and tumbling behavior with varying speeds and reorientation rates.

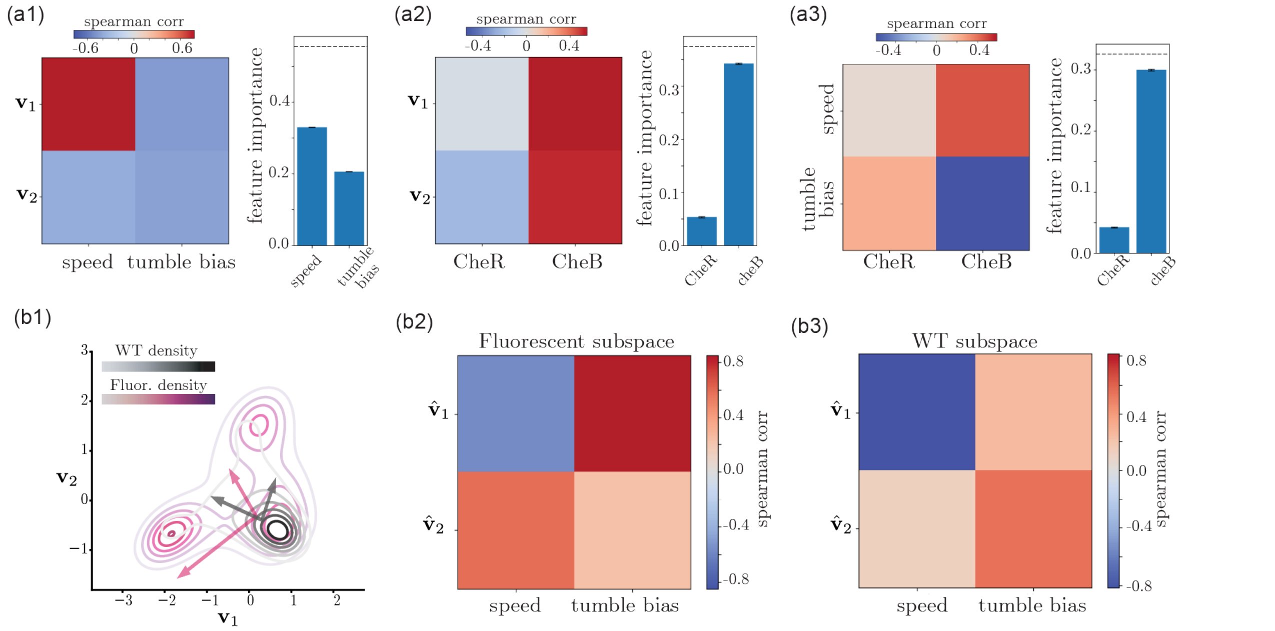


FIG. S4. **Phenotypic variation among wild-type and fluorescent bacteria is associated with variation in both run speed and tumble bias, and dominated by fluctuations in CheB.** (a1-left) Pairwise spearman correlation $\rho$ between the projection along the two main modes of variation $\mathbf{v}_1$ and $\mathbf{v}_2$, and the run speed and tumble bias of each individual. The first mode $\mathbf{v}_1$ is mostly correlated with run speeds, $\rho_{\mathbf{v}_1,\text{speed}} = 0.78\,(0.77, 0.79)$, and also with tumble bias, $\rho_{\mathbf{v}_1,\text{tumble bias}} = -0.46\,(-0.48, -0.44)$. The second mode $\mathbf{v}_2$ is nearly equally correlated with tumble bias and speed, $\rho_{\mathbf{v}_2,\text{speed}} = -0.39\,(-0.41, -0.36)$, $\rho_{\mathbf{v}_2,\text{tumble bias}} = -0.44\,(-0.46, -0.42)$. Unless otherwise noted, p-values are negligibly small. Each estimate of the correlation is accompanied by bootstrapped 95% confidence intervals. (a1-right) Importance of either speed or tumble bias in predicting the position of each individual along ($\mathbf{v}_1$,$\mathbf{v}_2$) using linear regression. We measure this by estimating the reduction in the coefficient of determination $R^2$ that results from shuffling either variable. Speed is more important than tumble bias in determining the position of each individual in the reconstructed phenotypic space. Error bars represent 95% bootstrapped confidence intervals over random reshuffles. The horizontal dashed line represents the $R^2$ of the unshuffled model. (a2-left) Same as (a1), but correlating $\mathbf{v}_1$ and $\mathbf{v}_2$ to CheR and CheB number in the mutant strains. Both modes are strongly correlated with CheB numbers, $\rho_{\mathbf{v}_1,\text{cheB}} = 0.55\,(0.52, 0.59)$, $\rho_{\mathbf{v}_2,\text{cheB}} = 0.52\,(0.48, 0.55)$, with $\mathbf{v}_2$ also correlated with CheR expression, $\rho_{\mathbf{v}_2,\text{cheR}} = -0.23\;(-0.28, -0.18)$, whereas $\mathbf{v}_1$ is uncorrelated with CheR, $\rho_{\mathbf{v}_1,\text{cheR}} = -0.03\,(-0.08, 0.03)$ (p-value=0.33). (a2-right) Same as (a1-right), but for the importance of either CheB or CheR in predicting the position of each individual along ($\mathbf{v}_1$,$\mathbf{v}_2$). The full model yields an $R^2 = 0.38\,(0.35, 0.40)$, with CheB fluctuations capturing most of the predictive power. (a3-left) Same as (a1), but correlating CheR and CheB numbers in the mutant strains to their run speed and tumble bias. Speed is uncorrelated to CheR numbers, $\rho_{\text{speed,cheR}} = -0.03\,(-0.02, 0.09)$ (p-value=0.23), but correlates with CheB, $\rho_{\text{speed,cheB}} = -0.46\,(0.42, 0.50)$. Tumble bias correlates most strongly with CheB, $\rho_{\text{tumble bias,cheB}} = -0.59\,(-0.62, 0.55)$, but also with CheR, $\rho_{\text{tumble bias,cheR}} = 0.22\,(0.18, 0.27)$. (a3-right) Same as (b1), but for the importance of CheB and CheR in predicting both the run speed and tumble bias of each individual. Once again, CheB is a major determinant of inter-individual variability. (b1) Probability distribution function (as a contour plot), and major axis of variation (arrows), for either the mutant data, or the wild-type bacteria. We find that the variation is more anisotropic in the mutant data when compared to wild-type data, and that the main axis of variation from either dataset are rotated by nearly 90° clockwise. (b2) Same as (a1-left), but estimating the Spearman correlation between the projection along the major axis of variation in the fluorescent dataset, and the speed and tumble bias. The main mode of variation in the fluorescent data $\hat{v}_1$ is mostly correlated with tumble bias, $\rho_{\hat{\mathbf{v}}_1,\text{tumble bias}} = 0.85\,(0.83, 0.86)$, but also speed, $\rho_{\hat{\mathbf{v}}_1,\text{speed}} = -0.57\,(-0.61, -0.52)$. The second mode is mostly correlated with speed, $\rho_{\hat{\mathbf{v}}_2,\text{speed}} = 0.59\,(0.55, 0.62)$, and less with tumble bias, $\rho_{\hat{\mathbf{v}}_2,\text{tumble bias}} = 0.23\,(0.18, 0.28)$. (b3) Same as (b2), but for the dominant modes of variation in the wild-type dataset. We find that $\mathbf{v}_1$ is mostly correlated with speed, $\rho_{\hat{\mathbf{v}}_1,\text{speed}} = -0.81\,(-0.83, -0.80)$, but also somewhat with tumble bias, $\rho_{\hat{\mathbf{v}}_1,\text{tumble bias}} = 0.26\,(0.23, 0.28)$. Conversely, $\mathbf{v}_2$ is mostly correlated with tumble bias, $\rho_{\hat{\mathbf{v}}_2,\text{tumble bias}} = 0.55\,(0.53, 0.57)$, and less with speed, $\rho_{\hat{\mathbf{v}}_2,\text{speed}} = 0.13\,(0.10, 0.16)$.

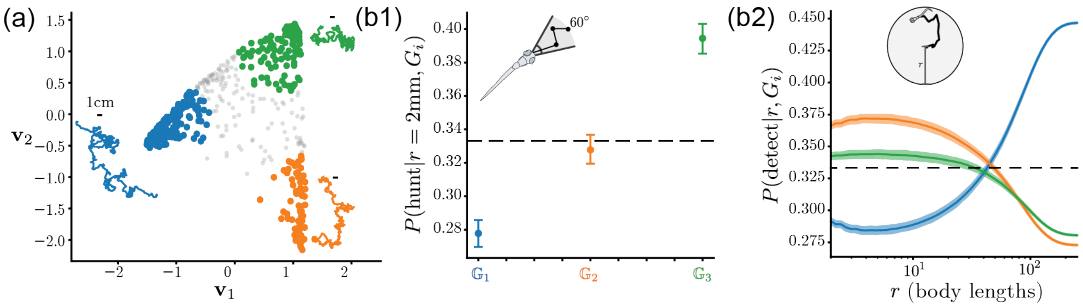


FIG. S5. **Analysis of the phenotypic groups from the larval zebrafish dataset.** (a) We utilize fuzzy c-means to define 3 groups at the extremes of the phenotypic space (see Methods). (b1) As in [16], we simulate boundaryless trajectories from the average transfer operator of the three groups. We calculate the probability of approaching a prey item successfully in each group by measuring the probability of resource collection in a 60° field of view from the start point within the first 3 bouts. The dotted line represents the average expected probability among 3 groups of 1/3. We find that $G_3$ is the best phenotype at hunting prey, matching the region with highest eye convergence in the phenotypic space, Fig. 5. (b2) We also calculate the probability of resource collection evenly distributed in circles of radius $r$ in simulated trajectories of 1000 bouts in duration. The bold black line represents the average expected behavior over 3 groups. We find that $G_1$ is the best at long range exploration, while $G_2$ and $G_3$ are good at local exploitation.

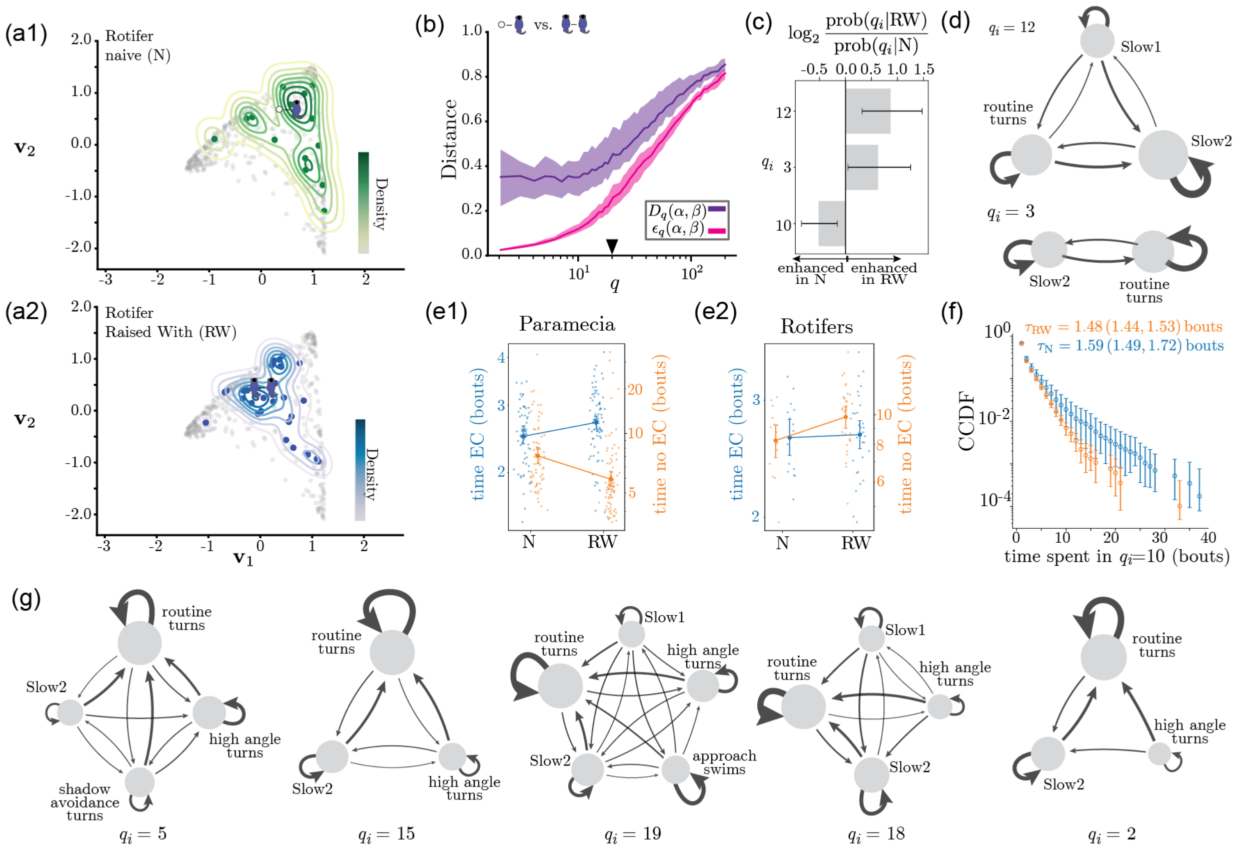


FIG. S6. **Details of the analysis of behavioral changes upon prior exposure in rotifers and paramecia.** (a) We compare naive (a1) vs experienced (a2) fish hunting rotifers. Prior exposures to rotifer shifts the distribution towards more exploratory phenotypes. (b) Distance $D_q$ and uncertainty $\varepsilon_q$ between naive and experienced fish hunting rotifers. While there is relevant information for a large number of of $q$, we compare conditions at $q = 20$ as in Fig. 6. (c) We report the log-fold change of the frequency of state usage at $q = 20$ for statistically relevant differences with p-value $< 0.05$ (two-sided permutation test) and $|\log_2 \frac{\text{prob}(q_i|\text{RW})}{\text{prob}(q_i|\text{N})}| > 0.5$. We find that raising fish with rotifers enhances the use of $q_i = 12$ and $q_i = 3$, while naive fish use more the $q_i = 10$, which was enhanced in the Paramecia RW condition. (d) We plot the transition matrix among bout types for $q_i = 12$ and $q_i = 3$, which corresponds to sequences of Slow 1, Slow2 and Routine Turns, characteristic of exploratory behavior. (e) Bout sequence duration with (blue) or without (orange) eyes converged in the conditions in which fish is either hunting paramecia (e1) or rotifers (e2). Small dots represent the geometric mean bout sequence durations of each fish; larger markers and error bars show the average and its 95% bootstrap confidence interval across fish. We find that fish raised with paramecia reduce the time spent with the eyes not converged (two-sided permutation test p-value $< 0.001$) and increase the time spent with eyes converged (two-sided permutation test p-value $< 0.01$). In contrast, when fish are raised with rotifers, we find that the time spent with eyes converged is unchanged (two-sided permutation test p-value $= 0.8$), but the time spent with the eyes not converged increases upon raising the fish with prey (two-sided permutation test p-value $= 0.028$). (f) Complementary Cumulative Distribution Function (CCDF) of the sequence length in state $q_i = 10$ for the conditions in which the fish is hunting rotifers. Unlike fish hunting paramecia, Fig. 6(e), we find that the sequence length in state $q_i = 10$ is reduced after raising the fish with rotifers: the geometric mean duration was $\tau_{\text{N}} = 1.59\,(1.49, 1.72)$ bouts for naive fish and $\tau_{\text{RW}} = 1.48\,(1.44, 1.53)$ bouts for fish raised with paramecia, where values in parentheses indicate the 95% confidence interval obtained by bootstrapping across fish. The difference between conditions was significant in a two-sided permutation test p-value $= 0.036$. The CCDF was obtained using all the available bout sequences, and error bars represent 95% confidence intervals bootstrapped across fish. (g) We plot the other transition matrices enhanced in the Paramecia Naive condition, Fig. 6(c). They mostly constitute more exploratory bout sequences, with Slow1, Slow2, and variable turns. We also find $q_i = 19$ with approach swims, which could represent a form of prey capture sequence used less after raising the fish with paramecia.

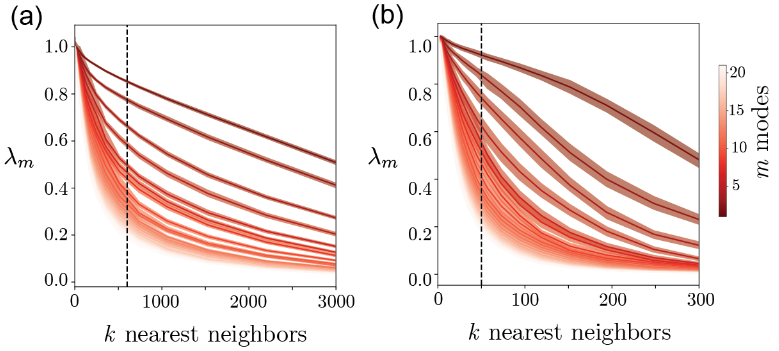


FIG. S7. **Selecting number of nearest neighbors $k$ for the self-tuning diffusion map for bacterial motility and larval zebrafish examples.** We calculate the eigenspectrum of the diffusion map for different values of $k$ for both systems. (a) Eigenvalues $\lambda_m$ for the bacterial locomotion phenotypic space showcasing the first $m$ eigenvalues of the system for multiple $k$. We select $k = 600$ neighbors, at a point where the dominant modes separate more cleanly from the bulk. (b) Eigenvalues $\lambda_m$ for the larval zebrafish phenotypic space showcasing the first $m$ eigenvalues of the system for multiple $k$. We select $k = 50$, for which the eigenvalue separation is clear.

### DIFFUSION MAPS CAPTURE THE INTRINSIC GEOMETRY OF PHENOTYPIC VARIATION

We consider that our time series observations, indexed by $\alpha$, result from a partial observation of an underlying Markovian stochastic process parameterized by $\theta^\alpha \in \mathbb{R}^N$ encoding underlying biological mechanisms. Changing in the underlying parameters map non-trivially to the emergent dynamics, and we use diffusion maps to probe the intrinsic geometry of the space of dynamics. Here, leveraging the connection between diffusion map eigenvectors and the Laplace-Beltrami operator [26], we show that the gradient of the diffusion map eigenvectors aligns locally with the dominant modes of the underlying metric tensor mapping variation in parameters to differences in dynamics.

We assume that in sufficiently local regimes the distance between observations $D$ admits a quadratic expansion,

$$D(\theta, \theta+\delta)^2 = \delta^\top g_D(\theta)\, \delta + o(\|\delta\|^2), \tag{S1}$$

This, in turn, induces a Riemannian metric tensor $g_D(\theta)$ on parameter space whose eigendirections define, locally in parameter space, which directions most impact the emergent dynamics. In addition, $g_D(\theta)$ defines a Laplace-Beltrami operator whose eigenfunctions capture the global intrinsic organization of the geometry induced by the distance between phenotypes, $D$. According to the Rayleigh-Ritz variational characterization of Laplace-Beltrami eigenfunctions [65, 66], the non-trivial eigenfunctions of the Laplace-Beltrami operator, $f$, minimize the Dirichlet energy,

$$\mathcal{E}[f] = \int (\nabla f)^\top g_D^{-1} (\nabla f)\, \mathrm{dvol}_g, \tag{S2}$$

subject to orthogonality constraints to lower modes. Consequently, low-frequency eigenfunctions vary preferentially along directions in parameter space where the induced geometry penalizes changes the least. This becomes clear when we project the gradient of the Laplace-Beltrami eigenfunctions locally in terms of the eigenspace of the underlying metric tensor $g_D(\theta)$. Let $g_D(\theta)e_a(\theta) = \Lambda_a(\theta)e_a(\theta)$, $\Lambda_a \geq \Lambda_{a+1} > \ldots > 0$, be the local eigensystem of the metric tensor $g_D(\theta)$. Writing, $\nabla f = \sum_a \gamma_a e_a$, the energy density becomes

$$(\nabla f)^\top g^{-1} (\nabla f) = \sum_a \frac{\gamma_a^2}{\Lambda_a}. \tag{S3}$$

Thus, in regions of parameter space where the metric $g_D(\theta)$ is strongly anisotropic, its dominant eigenvector $e_1(\theta)$ identifies the parameter direction along which the distance is locally most sensitive. Because the Dirichlet energy involves $g_D^{-1}$, gradients of low-frequency Laplace-Beltrami eigenfunctions can vary most cheaply along this same direction. We therefore expect the gradients of the leading diffusion-map eigenvectors to align locally with $e_1$.

We test this prediction in a cusp catastrophe model, Fig. S8, and illustrate how the result depends on the choice of distance $D$: different notions of distance between the stochastic dynamics induce different metric tensors and therefore identify different directions in parameter space as dynamically important.

We study a stochastic cusp catastrophe model,

$$\mathrm{d}X_t = -\partial_x V(X_t; \mu, c)\, \mathrm{d}t + \sqrt{2T_x}\, \mathrm{d}W_t, \text{where } V(x; \mu, c) = \frac{x^4}{4} - \mu\frac{x^2}{2} + cx, \tag{S4}$$

which exhibits either one or two fixed points depending on $\mu$ and $c$, and is either right or left biased depending on the sign of $c$,Fig. S8(a). We first consider a path-space distance, for which both the local Hellinger geometry and the associated path-space Fisher information rate can be determined analytically. Let $\mathbb{P}_\theta$ denote the path measure generated by the Langevin dynamics at parameter value $\theta$. Since two parameter values $\theta, \theta'$ differ only in drift, while sharing the same noise strength, Girsanov theory gives the local path-space Fisher information rate,

$$\frac{1}{T}\mathrm{KL}\left(\mathbb{P}_\theta \,\|\, \mathbb{P}_{\theta+\delta}\right) = \frac{1}{2}\delta^\top I_{\mathrm{path}}(\theta)\delta + o(\|\delta\|^2), \tag{S5}$$

where

$$I_{\mathrm{path}}(\theta)_{ij} = \frac{1}{2T_x}\mathbb{E}_{\pi_\theta}\left[\partial_{\theta_i}\partial_x V(X;\theta)\, \partial_{\theta_j}\partial_x V(X;\theta)\right].n \tag{S6}$$

For the quartic potential, we have,

$$I_{\mathrm{path}}(\mu, c) = \frac{1}{2T_x}\begin{pmatrix} \mathbb{E}_{\pi_{\mu,c}}[X^2] & -\mathbb{E}_{\pi_{\mu,c}}[X] \\ -\mathbb{E}_{\pi_{\mu,c}}[X] & 1 \end{pmatrix}, \tag{S7}$$

where $\mathbb{E}_{\pi_{\mu,c}}$ denotes expectation under the stationary density $\pi_{\mu,c} \propto e^{-V(x;\mu,c)/T_x}$. Although the Fisher tensor is naturally derived from the local expansion of the KL divergence, the KL divergence is asymmetric and is therefore not ideal as a distance for constructing diffusion-map kernels. We instead use the corresponding local Hellinger-rate distance, which is symmetric and induces the same local Fisher geometry up to a constant factor. For two parameter values $\theta_a = (\mu_a, c_a)$ and $\theta_b = (\mu_b, c_b)$, let $\theta_m = (\theta_a + \theta_b)/2$. We define the symmetric midpoint approximation

$$D_H^2(\theta_a, \theta_b) = \frac{1}{16T_x} \int \left[(\mu_b - \mu_a)x - (c_b - c_a)\right]^2 \pi_{\theta_m}(x)\, dx. \tag{S8}$$

This expression is the local quadratic approximation to the squared Hellinger distance per unit time between path measures. Indeed, for infinitesimal perturbations,

$$D_H^2(\theta, \theta + \delta) = \frac{1}{8}\delta^\top I_{\text{path}}(\theta)\delta + o(\|\delta\|^2). \tag{S9}$$

Thus, the path-space Hellinger-rate distance is Fisher-local: it has the same eigendirections as $I_{\text{path}}(\theta)$. Therefore, following the argument of Eqs. S1,S2,S3, we expect the diffusion map eigenvectors computed from Hellinger-rate distances to vary along the stiff directions of the path-space Fisher information matrix. Fig. S8(b1) tests the resulting prediction. At each point in parameter space, we compare the gradient of a diffusion-map eigenvector with the dominant eigenvector of the local Fisher metric, and quantify their alignment by

$$A_k = \sum_i \omega_i \frac{|\nabla_\theta \mathbf{v}_k(\theta_i) \cdot e_1(\theta_i)|}{\|\nabla_\theta \mathbf{v}_k(\theta_i)\|}, \tag{S10}$$

where $\omega_i \propto \frac{\Lambda_1(\theta_i)}{\Lambda_2(\theta_i)}$ emphasizes regions in which the metric is strongly anisotropic. The gradients of the leading diffusion coordinates show strong local alignment with the dominant Fisher direction, as predicted by the geometric argument above. The stiff mode of the path-space Fisher information rate is aligned parallel to the bifurcation boundary, highlighting how the Hellinger distance between paths is most sensitive to infinitesimal changes in parameters integrated over the entire domain: the stiff mode in this case seems to vary with the stability of the most stable minimum of the potential, Fig. S8(a).

We next showcase how different distances highlight different properties of the dynamics, while the geometric argument holds. For example, we consider the Hellinger distance among 2-state Markov models, which compresses the finescale dynamics into discrete labels depending on the sign of $x$. In this case, we find that the stiff Fisher mode is orthogonal to the bifurcation boundary, highlighting how at the level of a 2-state description of the system this is the direction that most changes the transition dynamics at the longest timescales, Fig. S8(b2). Accordingly, the diffusion map eigenmodes vary with $\mu$ and $c$ but along directions that directly capture the transition between bistable and monostable solutions, and their gradients align with the Fisher information eigenvectors.

For completeness, we also assess the embedding obtained using our multiscale dissimilarity, Eq. 3. Despite being agnostic about the correct coarse-graining scale to compare the dynamics, our approach recovers the alignment of the diffusion map eigenvectors with the bifurcation structure and the asymmetry of the potential landscape. In addition, we find a very good agreement between the gradient of the diffusion map eigenvectors and the Fisher information metric induced by the Hellinger distance among 2-state Markov chains, indicating that our multiscale metric preserves the most relevant long-lived structure in the dynamics in an unbiased way.

### COMPARING THE SPECTRA OF TRANSFER OPERATORS

Beyond the path-space Fisher information rate, another common approach for comparing dynamical systems is to compare the spectra of their transfer operators [47–50]. In particular, the optimal transport distances between their singular-value spectra provides a compact measure of changes in dynamical structure, including changes in the number and stability of fixed points. Our framework provides a convenient way to compute such distances because the construction of our coarse-grained states already relies on Ulam-Galerkin approximations of Perron-Frobenius operators (Methods). We can therefore directly compare the singular value spectra of these operators using a Wasserstein distance.

The Lorenz system [67] illustrates the strength of this spectral comparison, Fig. S9(a,b,c). As parameters are varied, the Wasserstein distance between transfer operator spectra distinguishes qualitatively different dynamical regimes, including fixed points, limit cycles, and chaotic dynamics, allowing these phases to be recovered in the corresponding diffusion map embedding.

The stochastic cusp model, however, exposes an important limitation of this representation, Fig. S9(d). Applied to Eq. S4, the resulting embedding is primarily sensitive to changes in $\mu$, but does not distinguish $\theta = (\mu, c)$ from $\theta = (\mu, -c)$. Changing the sign of $c$ reflects the potential about $x = 0$. The two systems therefore have the same dynamical organization, including the same number and stability of fixed points, while exchanging the identities of the left- and right-biased states. Their transfer-operator spectra consequently provide no information about which of these states is being modulated. For some applications this invariance is desirable: it identifies systems with the same underlying dynamical structure irrespective of the labeling or location of their states. For dynamical phenotypes, however, the identity of the affected state can itself be biologically meaningful. In comparison, our multiscale dissimilarity retains sensitivity to changes in metastable structure while also keeping track of which dynamical states are changing.

### DETAILS OF NUMERICAL ANALYSIS

#### Stochastic cusp catastrophe

Instead of making simulations, we here directly approximate the infinitesimal generator of the Fokker-Planck dynamics using the Chang-Cooper method [68], discretizing the domain $x \in [-\sqrt{\mu_{\max}} - 4\sqrt{T_x}, \sqrt{\mu_{\max}} + 4\sqrt{T_x}]$ uniformly into 1000 points, where $\mu_{\max}$ is the largest sampled value of $\mu$. The value of $\mu$ was sampled between 0 and 1.4, with a uniform spacing of $\Delta\mu = 0.05$, whereas $c$ values were sampled between -0.4 and 0.4, with a uniform spacing of $\Delta c = 0.025$.

| dynamical regime | $r$ |
|---|---|
| stable fixed point | 20 |
| periodic $[1,1]$ | 220 ; 250 |
| periodic $[1,2,2]$ | 100.5 |
| periodic $[1,2,2]^2$ | 99.7 |
| periodic $[1,2,2]^4$ | 99.6 |
| periodic $[1,2,2]^8$ | 99.537 |
| periodic $[1,1,2,2]$ | 152 , 165 |
| periodic [1, 1, 2, 2, 1, 2 | 126.5 |
| intermittent chaos | 100.93 |
| chaos | 28 ; 45 ; 60 ; 75 ; 99.4 ; 115 ; 140 ; 195 |

TABLE I. Sampled dynamical regimes of the Lorenz system across $r$.

#### Lorenz system

We simulate time series from the equations of motion,

$$\begin{cases} \dot{x} &= \sigma(x - y) \\ \dot{y} &= (r - z)x - y \\ \dot{z} &= xy - \beta z, \end{cases} \tag{S11}$$

with $\sigma = 10$, $\beta = 8/3$ and varying $r$ sampled to capture different dynamical regimes [69] as in [49] (see Table I). We sampled from each value of $r$ 10 times, using the `scipy.odeint` package [70] to simulate for $T_{\text{expt}} = 1000\,\text{s}$ with a sampling time of $\delta t = 0.01\,\text{s}$, discarding the first 1000 s as transient dynamics. We then approximated the Perron-Frobenius operator as in [13] by partitioning the state space into $N = 1000$ partitions and using a transition time of $\tau = 0.1\,\text{s}$, obtaining a $N \times N$ transition operator $P_{ij}$ for each simulated timeseries. Finally, we estimated the Wasserstein distance between the singular values $\{\sigma_i\}$ of each $P_{ij}$, Fig. S9(a), and used self-tuning diffusion maps with a bandwidth inferred from 30 nearest neighbors to reconstruct the phenotypic space in Figs. S9(b,c). We trained an L1-regularized multinomial logistic regression classifier to predict the dynamical regimes from the diffusion map coordinates (keeping 20 dimensions). The classifier performance was assessed using repeated stratified

5-fold cross-validation with 100 repeats, and the confusion matrix shown in Fig. S9(b2) is averaged across all 500 splits. We used the Lyapunov spectrum to access how the diffusion map dimensions resolve different properties of the dynamics: unpredictability (as measured through the largest Lyapunov exponent), and dimensionality (as measured through the Kaplan-Yorke dimension [71]). Lyapunov spectra were estimated using a continuous Gram–Schmidt method, integrating the analytical Jacobian over 1000 s. For each parameter $r$, we repeated the calculation 5 times and averaged the resulting spectra over realizations.

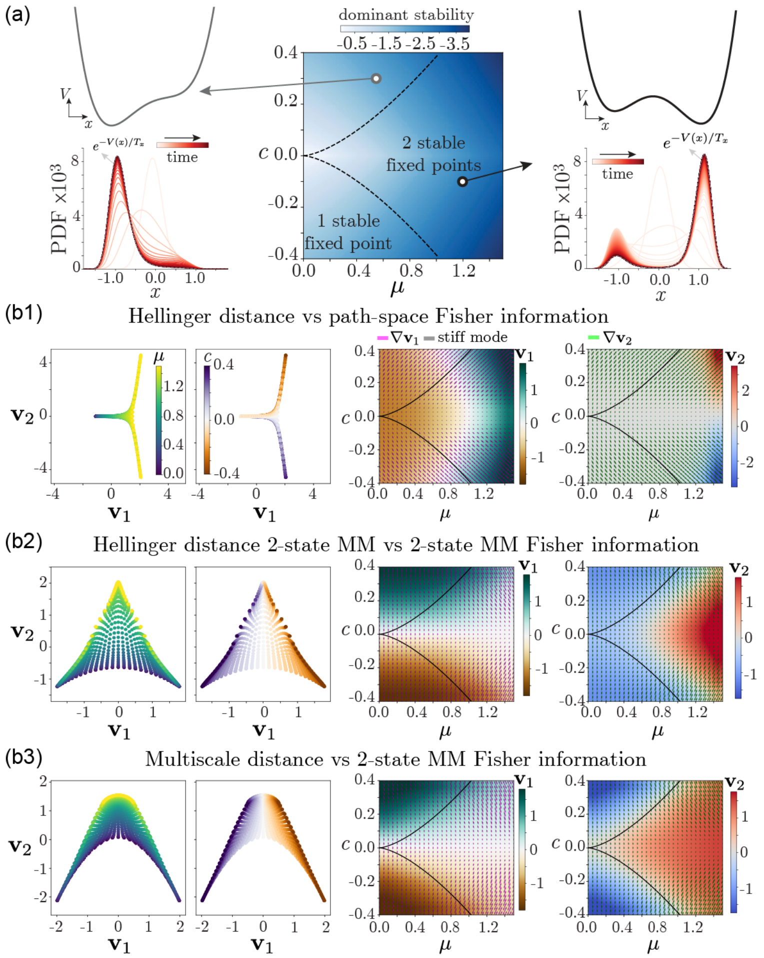


FIG. S8. **Relationship between diffusion map eigenvectors and the stiff directions of the induced metric mapping parameters to emergent dynamics.** (a) - We consider overdamped Langevin dynamics in a quartic potential. In the center, we plot a phase diagram as a function of $\theta = (\mu, c)$, separating the regime with 2 stable fixed points from the regime with 1 stable fixed point at boundary $(c/2)^2 = (\mu/3)^3$ (black dashed line). We color code the phase diagram by the curvature of the most stable fixed point. We also show examples of the potential landscape in the monostable (top) and bistable (bottom) regimes, as well as an example simulation of the density dynamics. (b) - We use diffusion maps to reconstruct the low-wavelength modes of the underlying Laplace-Beltrami operator, and showcase how the gradient of the diffusion map eigenvectors aligns with the stiff directions of the underlying metric. In each panel, we plot on the left the inferred diffusion map embedding, and on the right the parameter space color coded by each diffusion map eigenvectors, as well as the gradient of each eigenvector ($\nabla \mathbf{v}_1$ in pink and $\nabla \mathbf{v}_2$ in green) and the stiff mode of the induced metric tensor (grey). (b1) - Hellinger distance between paths, estimated using Eq. S8, compared against the path-space Fisher information matrix. We find that $\mathbf{v}_1$ is mostly associated with changes in $\mu$, and $\mathbf{v}_2$ is mostly associated with changes in $c$ (left). The direction of change of the diffusion map eigenvectors matches the local stiff directions of the path-space Fisher information matrix, with an alignment score of $A_1 = 0.971\,(0.967, 0.976)$ for $\mathbf{v}_1$ and $A_2 = 0.991\,(0.990, 0.992)$ for $\mathbf{v}_2$. Notice that the stiff mode of the path-space Fisher information aligns with the change in stability of the most stable fixed point (shown in a). (b2) - Same as (b1), but now comparing the Hellinger distance between 2-state Markov models with the corresponding Fisher information matrix. We coarse-grain the dynamics into two states along $x = 0$. We find that $\mathbf{v}_1$ mostly varies with $c$ whereas $\mathbf{v}_2$ mostly varies with $\mu$ (left). The gradient of the diffusion map agrees with the stiff mode of the underlying Fisher information matrix ($A_1 = 0.9992\,(0.9991, 0.9993)$, $A_2 = 0.95\,(0.94, 0.96)$). The dominant modes are orthogonal to the bifurcation boundary, separating monostable from bistable solutions. (b3) - Same as (b2), but now using the multiscale total variation distance introduced in the main text. The resulting embedding recovers the asymmetry of the potential landscape along $\mathbf{v}_1$, and the transition between bistable and monostable regimes along $\mathbf{v}_2$. The gradient of the diffusion map eigenvectors still agrees with the stiff modes of the Fisher information matrix computed from the Hellinger distance between 2-state Markov models ($A_1 = 0.979\,(0.976, 0.982)$, $A_2 = 0.938\,(0.930, 0.945)$).

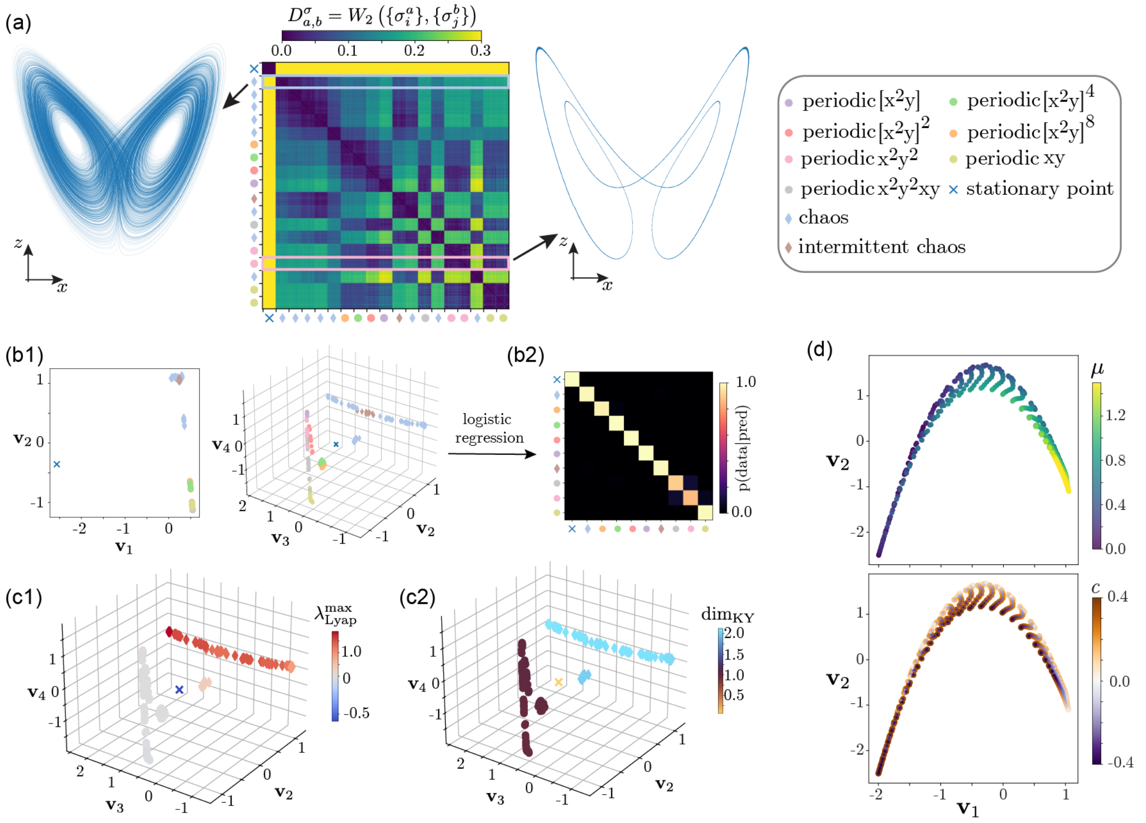


FIG. S9. **Comparing dynamics using the optimal transport distance between the singular values of the inferred transfer operators.** (a) Our approach provides an efficient approximation of Perron-Frobenius operators from time series data. We here showcase how the optimal transport distance among their singular values enables the recovery of different dynamical regimes in the Lorenz system [67], Eq. S11. We set $b = 8/3$, $\sigma = 10$ and vary $r$ to sample different dynamical regimes. We simulate 10 time series from different initial conditions for each value of $r$, and show example simulations (left: chaos $r = 28$, right: periodic solution $r = 165$). We estimated the Wasserstein distance between the singular values of an inferred Perron-Frobenius operators, $D^{\sigma}_{a,b} = W_2(\{\sigma^a_i\},\{\sigma^b_j\})$ (center). The tick marks represent different values of $r$. (b) Diffusion map embedding of the Wasserstein distance matrix. (b1) The first dimension $\mathbf{v}_1$ isolates out the stable fixed point solution from everything else (left); $\mathbf{v}_2$ captures the difference between limit cycle and chaotic solutions, $\mathbf{v}_3$ differences among chaotic solutions, and $\mathbf{v}_4$ differences among periodic solutions. (b2) We use a regularized logistic regression to classify the points in the diffusion map embedding into their respective dynamical regimes, resulting in classification accuracy of $0.97\pm0.04$ (see Section ). We plot the average confusion matrix obtained from repeated stratified 5-fold cross-validation (100 repetitions). (c) We examine how the different diffusion map dimensions map onto properties of the Lyapunov spectrum. (c1) We color code the diffusion map embedding by the largest Lyapunov exponents, showcasing how $\mathbf{v}_2$ distinguishes chaotic from non-chaotic dynamics, while $\mathbf{v}_3$ subsequently captures the degree of chaos. (c2) We color code the diffusion map embedding by the Kaplan-Yorke dimension [71], showcasing the difference between fixed point solutions (which lie along negative $\mathbf{v}_1$, visible in panel b1), limit cycle solutions and fractal chaotic solutions. (d) We reproduce the results on the cusp catastrophe example Fig. S8, but using a Wasserstein distance between singular values of $\mathcal{L}_\theta$. We approximate $\mathcal{L}_\theta$ using the Chang-Cooper method [68] for each $\theta = (\mu, c)$, and compute the singular values of $\mathcal{P}_\tau = e^{\mathcal{L}\tau}$ with $\tau = 0.1\,$s. The diffusion map embedding constructed with a self-tuning kernel (with 4 nearest neighbors), shows that $\mathbf{v}_1$ mostly aligns with variation in $\mu$, while $\mathbf{v}_2$ can capture the overall asymmetry of the potential, but cannot distinguish $\theta = (\mu, c)$ from $\theta = (\mu, -c)$ because these are topologically conjugate.
